# miRDriver: A Tool to Infer Copy Number Derived miRNA-Gene Networks in Cancer

**DOI:** 10.1101/652156

**Authors:** Banabithi Bose, Serdar Bozdag

**Author notes:** Permission to make digital or hard copies of part or all of this work for personal or classroom use is granted without fee provided that copies are not made or distributed for profit or commercial advantage and that copies bear this notice and the full citation on the first page. Copyrights for third-party components of this work must be honored. For all other uses, contact the owner/author(s).

## Abstract

Copy number aberration events such as amplifications and deletions in chromosomal regions are prevalent in cancer patients. Frequently aberrated copy number regions include regulators such as microRNAs (miRNAs), which regulate downstream target genes that involve in the important biological processes in tumorigenesis and proliferation. Many previous studies explored the miRNA-gene interaction networks but copy number-derived miRNA regulations are limited. Identifying copy number-derived miRNA-target gene regulatory interactions in cancer could shed some light on biological mechanisms in tumor initiation and progression. In the present study, we developed a computational pipeline, called miRDriver which is based on the hypothesis that copy number data from cancer patients can be utilized to discover driver miRNAs of cancer. miRDriver integrates copy number aberration, DNA methylation, gene and miRNA expression datasets to compute copy number-derived miRNA-gene interactions in cancer. We tested miRDriver on breast cancer and ovarian cancer data from the Cancer Genome Atlas (TCGA) database. miRDriver discovered some of the known miRNAs, such as miR-125b, mir-320d, let-7g, and miR-21, which are known to be in copy number aberrated regions in breast cancer. We also discovered some potentially novel miRNA-gene interactions. Also, several miRNAs such as miR-127, miR-139 and let-7b were found to be associated with tumor survival and progression based on Cox proportional hazard model. We compared the enrichment of known miRNA-gene interactions computed by miRDriver with the enrichment of interactions computed by the state-of-the-art methods and miRDriver outperformed all the other methods.

**CCS CONCEPTS:**

- Bioinformatics
- Computational Genomics
- Biological Networks

## 1 Introduction

Cancers, like many other diseases, can be defined as a disease of altered gene expression due to dysregulation at the transcriptional and post-transcriptional layers [1–2]. microRNAs (miRNAs) are short non-coding RNAs that found to regulate gene expression post-transcriptionally [3]. A single miRNA can corelate hundreds of genes and can influence the tumor suppressor genes and oncogenes [4].

miRNA dysregulation has been related to several types of cancer [5]. Many studies discovered the affiliation of miRNAs to the cancers driving genes in tumor initiation and progression. For example, miR.125b, miR.145, miR.21 and miR.155 are known to be involved in breast cancer [6] and, miR.27b and miR.181d are known to be involved in ovarian cancer [7–8].

Efforts were dedicated in integrating and analyzing high-throughput expression data to find cancer specific miRNA-gene interaction networks [9–10]. Mutual information-based methods such as AraCNe [11] and ProMISe [12] to infer gene regulatory networks received popularity. To explore the causal relationship among miRNA and gene (*i.e*. direct or indirect effects of miRNA on genes), invariant causal prediction (ICP) methods like Hidden-ICP, ICP-PAM50 emerged [13]. Other casual inference-based methods such as idaFast [14] and jointIDA [15] were also developed. These techniques were implemented using expression profiles of miRNAs and genes.

Several studies also explored the integration of multidimensional omics data in cancer [16] such as copy number alteration (CNA), and gene expression to infer gene regulatory interactions based on the premise that CNA regions could harbor driver regulatory genes in cancer [17–18]. Akavia et al., 2010 developed a Bayesian network-based computational tool, CONEXIC that integrates copy number and gene expression data for detecting aberrations in cancer progression. A few studies (Setty et al., 2010; Li et al., 2014), applied regression-based approach to integrate copy number, DNA methylation, and miRNA expression to predict gene expression changes in terms of transcription factors (TFs) and miRNA expression. In their efforts of integrating CNA, miRNA and gene expression, these studies used miRNAs from all genomic regions.

CNA regions have been reported to harbor key miRNAs, too [19]. Calin et al. [20] found that miR.15a and miR.16a were located in chromosome 13q14.3, which is frequently deleted in B cell chronic lymphocytic leukemias. Zang et al. [21] studied 283 miRNAs associated with breast cancers and confirmed that 72.8 % of miRNAs are in CNA regions.

Despite the existence of potential driver miRNAs in CNA regions, to the best of our knowledge, there is no computational method that mine these areas to identify key miRNAs and their potential targets. To address this gap, in the current study, we specifically considered the CNA regions to infer CNA-derived miRNA-gene interactions in cancer. We hypothesized that CNA regions may host driver miRNAs and these miRNAs could regulate key biological processes in cancer by targeting some downstream genes. We developed a computational pipeline called miRDriver, which integrates gene and miRNA expression, copy number alteration, DNA methylation and TF-gene interaction information to infer copy number-derived miRNA-gene networks in cancer. In miRDriver, we retrieved frequently aberrated copy number regions among cancer patients using GISTIC [22–23]. For each region, we computed differentially expressed genes (DE) between frequently aberrated patients and not-frequently aberrated patients, and applied a LASSO-based [24] regression model to select miRNAs which could potentially regulate DE genes’ expression.

We assessed miRDriver using breast cancer and ovarian cancer data from TCGA [25] database. miRDriver outperformed AraCNe, ProMISe, Hidden-ICP, ICP-PAM50, IDAfast and jointIDA in retrieving significantly enriched miRNA-gene interactions with the known miRNA-gene interaction databases. We observed that a higher number of selected prognostic miRNAs in frequently aberrated patients than in not-frequently aberrated cancer patients. Furthermore, the miRNAs selected by miRDriver were found to be enriched in known cancer-related miRNAs and our selected genes were found to be enriched in many cancer-related pathways.

## 2 Materials and methods

### 2.1 miRDriver

We developed miRDriver, a computational pipeline to infer copy number aberration-based miRNA-gene interactions in cancer. miRDriver integrates gene expression, copy number alteration, DNA methylation, transcription factor-gene interaction and miRNA expression data. miRDriver has four main computational steps (Fig.1). In the first step, we employed GISTIC tool to find frequently aberrated chromosomal regions among cancer patients and called these regions as GISTIC regions. In the second step, for each GISTIC region we computed differentially expressed (DE) genes between frequently aberrated and non-aberrated patient groups. In the third step, we retrieved DE genes and miRNAs that reside in aberrated regions (*i.e. cis* genes and *cis* miRNAs) and retrieved DE genes that are outside of aberrated regions (*i.e. trans* genes). In the last step, we employed a LASSO-based regression model to compute miRNA regulators of the *trans* genes. In what follows, we describe the four steps of miRDriver in details. The entire pipeline of miRDriver is illustrated in Fig.1. The source code for miRDriver is available at https://github.com/bozdaglab/miRDriver under MIT license. All the supplementary files referenced in this manuscript can be accessed on the same page.

**Figure 1:**
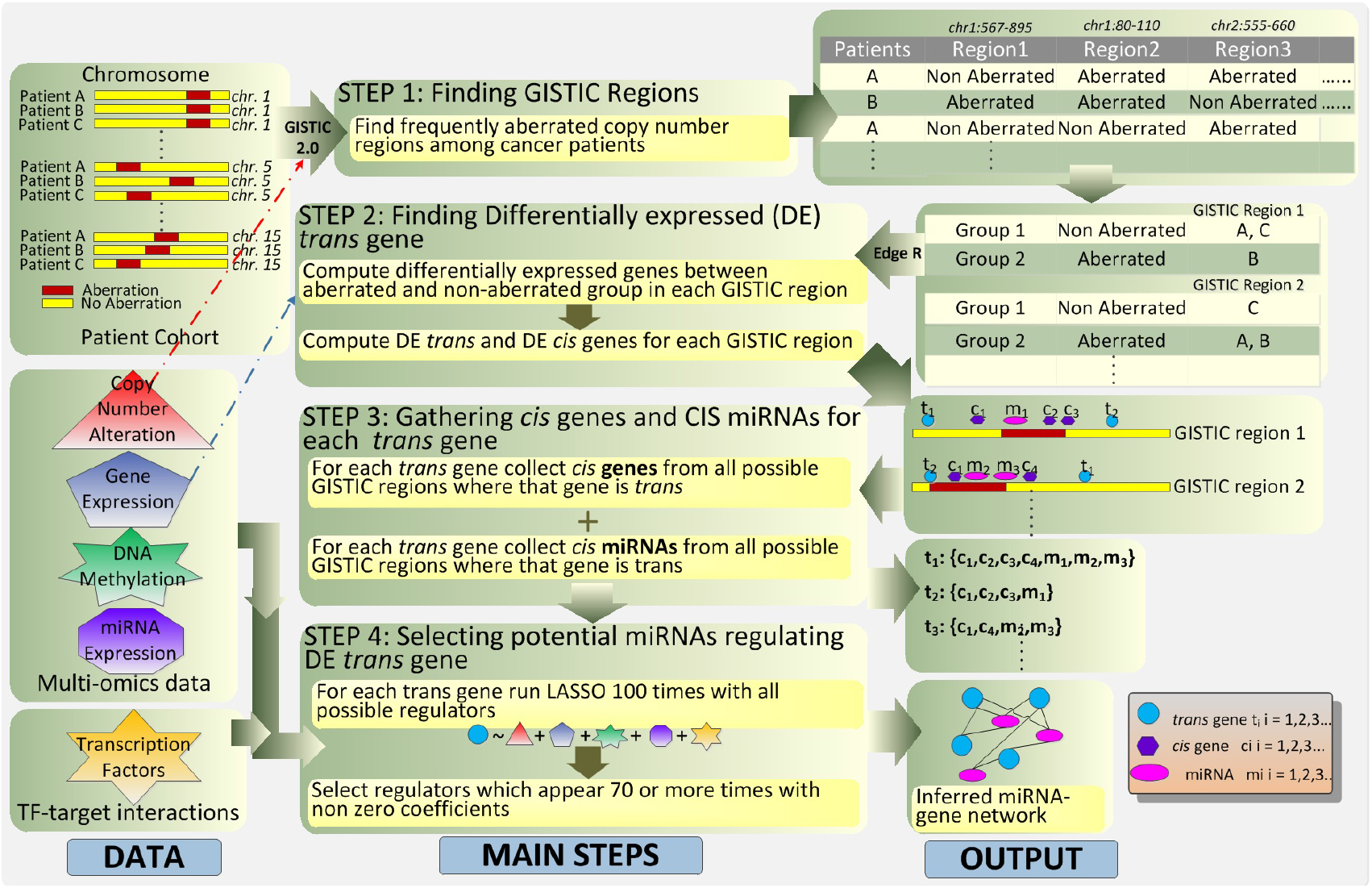
Flowchart of the miRDriver pipeline.

#### 2.1.1 Finding GISTIC regions

Our pipeline was developed based on the assumption that chromosomal regions that undergo CNA events among cancer patients could harbor driver miRNAs and these miRNAs could regulate downstream target genes in cancer. We used GISTIC 2.0 to identify chromosomal regions that were frequently aberrated within the patient cohort. We used a high confidence interval level of 0.90 to calculate frequently aberrated chromosomal regions (hereafter GISTIC regions). GISTIC regions with a log_2_ ratio above 0.1 were considered amplified and GISTIC regions with a log_2_ ratio below −0.1 were considered deleted.

#### 2.1.2 Finding DE genes

To identify transcriptional effect that is potentially due to a CNA event, we computed DE genes for each GISTIC region. For each GISTIC region, we divided cancer patients into two groups, frequently aberrated group and not-frequently aberrated group. We used *edgeR* [26] package in R to compute DE genes among these two groups (absolute log fold change (logFC) ≥ 1, adjusted p-value ≤ 0.05). We annotated DE genes that were located inside the GISTIC region as *cis* genes and DE genes outside of the GISTIC region were considered as *trans* gene.

#### 2.1.3 Gathering cis genes and cis miRNAs

In this step, we retrieved miRNAs located in each GISTIC region (*i.e. cis* miRNAs). The Cancer Genome Atlas (TCGA) [25] database for miRNA Gene Quantification data and Isoform data containing chromosomal regions were used to retrieve miRNAs reside within each GISTIC region using *bedr* package in R. Then, for each *trans* gene, we collected all *cis* genes and *cis* miRNAs from all the GISTIC regions where gene was appeared as DE *trans* gene. At the end of this step we had all possible *trans-cis* combinations for the entire patient cohort. We considered only those *trans* genes that had at least one *cis* miRNA predictor.

#### 2.1.4 Selection of miRNA regulators for each trans gene

For each DE *trans* gene miRDriver predicted the associated *cis* miRNAs that influenced the *trans* gene’s expression variation. Because *trans* gene’s expression can be influenced by other potential regulators such as *cis* genes in the same GISTIC region(s), TFs that are known to target this *trans* gene, and CNA and DNA methylation signals of this *trans* gene, we applied a LASSO-based predictor selection method with all these potential predictors as independent variables and the trans gene’s expression as response variable. We applied 10-fold cross validation to find the optimal regularization hyperparameter λ that provided the simplest model such that its cross-validation error was the minimum cross-validation error. For each *trans* gene, out of all its candidate predictors (independent variables), LASSO regression selected a set of non-zero coefficient predictors. We employed R package *glmnet* [27] to perform LASSO. Since the independent variables selected by LASSO have been shown to be inconsistent especially when sample size gets large [28], for each *trans* gene we ran LASSO 100 times. Only the *cis* miRNAs whose coefficients appeared to be non-zero at least 70 times were selected as potential regulator miRNAs.

### 2.2 Datasets

#### 2.2.1 Datasets to run miRDriver

We assessed miRDriver on breast and ovarian cancer datasets from TCGA. We used TCGABiolinks [29] to download the genomic data of cancer patients from TCGA. We retrieved gene expression quantification data for raw count mRNA (Illumina HiSeq), miRNA-gene quantification expression with file type hg19.mirbase20.mirna and miRNA isoform gene quantification data with file type hg19.mirbase20.isoform from the legacy data.

For mRNA data (*i.e*. RNA sequencing data), lowly expressed genes were eliminated and rest of the mRNA expression were converted to RPKM (Reads Per Kilobase of transcript, per Million mapped reads) values. We used the miRNAs that have ≥ 0.01 RPM (Reads per million mapped reads) value across ≥ 30% of the entire cohort.

We retrieved masked copy number variation (Affymetrix SNP Array 6.0) and computed the gene-centric copy number value compatible with hg38 using R Bioconductor package *CNTools* [30].

We downloaded DNA methylation data from Infinium HumanMethylation27 Bead-Chip (27K) and Infinium HumanMethylation450 Bead-Chip (450K) platforms. Gene-specific beta values were calculated separately for both platforms. For the 450K platform, average beta value for promoter-specific probes were considered due to their role in transcriptional silencing [31]. Given lower coverage in the 27K platform, we utilized all the probes. In this case, we set the DNA methylation of a gene as the average of beta values of all its probes.

To incorporate TF-gene interactions in the LASSO step (Section 2.1.3), we downloaded experimentally-validated TF-gene interactions from TRED [32] and TRRUST [33] databases.

#### 2.2.2 Datasets to evaluate miRDriver’s results

To evaluate the selected miRNA-gene interactions by miRDriver, we use the list of experimentally-verified miRNA-gene interactions used in [34] as our ground truth. Since miRDriver could identify indirect downstream targets (i.e., target of a direct target) as well as direct targets, we included potential indirect targets to the ground truth dataset. Specifically, for each miRNA-gene interaction where gene is a known TF, we included the experimentally-validated targets of this TF as the indirect target of this miRNA. We downloaded experimentally-validated TF-target regulatory datasets from TRED [32] and TRRUST [33] databases. For clinical data we used TCGA Clinical Data Resource [35] with clinical outcome endpoints for BRCA and OV.

To evaluate if the miRNAs selected by miRDriver were enriched in cancer-related miRNAs, we downloaded a list of known cancer related miRNAs from oncomiRDB database [36]. Each miRNA listed in oncomiRDB is involved in at least one cancer-related phenotype or cellular process. We harmonized the names of oncomiRDB miRNAs with reference to miRBase [37] database.

## 3 Results

We developed a computational pipeline called miRDriver which integrates CNA, DNA methylation, TFs-gene interactions, gene and miRNA expression datasets to compute copy number-derived miRNA-gene interactions in cancer.

We assessed miRDriver using breast and ovarian cancer datasets from TCGA. We downloaded RNA sequencing data of primary tumors in 1097 and 586, breast invasive carcinoma (BRCA) and ovarian serous cystadenocarcinoma (OV) patients, respectively. We used copy number segmentation data from TCGA to find frequently aberrated chromosomal regions, namely, GISTIC regions using GISTIC 2.0 (see Section 2.1.1). We merged overlapping GISTIC regions using R Bioconductor package *bedr* and considered 66 and 64 GISTIC regions for BRCA and OV, respectively. Among these regions, 43 and 56 GISTIC regions harbor miRNAs in BRCA and OV, respectively. We used R package *edgeR* to compute DE *trans* and *cis* genes for each GISTIC region (see Section 2.1.2). Using R package *bedr*, we retrieved miRNAs from each GISTIC region (see Section *2.1.3*). Altogether, 255 *cis* miRNAs for BRCA and 301 *cis* miRNAs for OV were used in miRDriver. We provided the count of DE c*is* genes, *cis* miRNAs and DE *trans* gene for GISTIC regions in each cancer type in Supplemental Table 1. We employed a LASSO-based regression with *trans* gene’s expression as response variable and expression of *cis* miRNAs as independent variables. In order to account for other potential regulators of *trans* gene’s expression, we also included expression of *cis* genes, copy number values of the *trans* gene, DNA methylation beta values of the *trans* gene and expression of TFs targeting that *trans* gene as independent variables in LASSO. We employed 10,494 *trans* genes for BRCA and 3,117 *trans* genes for OV (see Section 2.1.4). For each *trans* gene we ran LASSO 100 times. After LASSO, there were 1,858 and 1,347 selected miRNA-gene interactions for BRCA and OV, respectively. Among them BRCA had 187 miRNAs and 1,114 genes and OV had 147 miRNAs and 548 genes. The selected miRNA-gene interactions by miRDriver for each cancer type are listed in Supplemental Table 2.

## 4 Evaluation of miRDriver’s results

We performed several tests to assess the results of miRDriver. To test accuracy of the miRNA-gene interactions inferred by miRDriver, we performed a hypergeometric test between the computed targets of each miRNA and their known targets in our ground truth dataset (see Section 2.2.2). We also compared miRDriver with some of the existing methods and observed that miRDriver outperformed them in selecting significantly enriched mRNA-gene interactions. Also, we checked miRDriver’s performance in different subsets of breast cancer patients and found that there was significant overlap in miRDriver’s results on random subsets of the data. We also performed enrichment test for the selected miRNAs with the experimentally-validated oncogenic miRNA in oncomiRDB. The selected miRNAs by miRDriver were significantly enriched in the oncogenic miRNAs.

In the following sections we presented the details of miRDriver evaluation.

### 4.1 Computed miRNA-gene interactions were enriched in the known miRNA-target interactions

To check if the miRNA-gene interactions computed by miRDriver were significantly enriched in the known miRNA-gene interactions, we performed a hypergeometric test (see Section 2.2.2). We considered only those miRNAs that had at least one known target in the ground truth data for the hypergeometric test. For breast cancer 59 miRNAs and for ovarian cancer 27 miRNAs were used in the hypergeometric test. For breast cancer 63% predicted miRNAs and for ovarian cancer 89% miRNAs showed significant enrichment (p-value ≤ 0.05). The hypergeometric test result for a few miRNAs is provided in Table 1 with the count of computed targets and p-value for each miRNA. The entire list of selected miRNAs with hypergeometric result is listed in Supplemental Table 3.

**Table 1:** Enrichment results of computed miRNA-gene interactions in the known miRNA-gene interaction in BRCA and OV. P-value column shows the hypergeometric p-value.

| Cancer type | miRNA | No. of computed targets | p-value |
| --- | --- | --- | --- |
| BRCA | miR.582 | 162 | $9.6e^{-10}$ |
| | miR.339 | 75 | $3.8e^{-2}$ |
| | miR.330 | 42 | $6.3e^{-6}$ |
| | miR.744 | 41 | $3.1e^{-2}$ |
| | miR.1224 | 24 | $4.7e^{-3}$ |
| OV | let.7b | 35 | $1.7e^{-2}$ |
| | miR.181c | 23 | $1.4e^{-1}$ |
| | miR.33a | 14 | $3.2e^{-3}$ |
| | miR.27a | 12 | $3.1e^{-3}$ |
| | miR.3619 | 11 | $3.7e^{-2}$ |

### 4.2 miRDriver outperformed six existing methods for computing miRNA-gene interactions

We compared the accuracy of miRNA-gene interactions computed by miRDriver with six other popular methods namely, AraCNe, hiddenICP, ProMIse, idaFast, jointIDA and icpPAM50 using BRCA data. For AraCNe, ProMIse and hiddenICP we used 6504 input genes and 255 miRNAs, the same inputs we used in miRDriver. We applied different thresholds to select reported miRNA-gene interactions based on reported scores to get highly confident interactions for our comparison.

We used the AraCNe implementation in the R package *minet* [38] and computed 273 miRNA-gene interactions with non-zero scores and used all of them. For the other methods, we used R package *miRLAB* [39]. After applying ProMIse method, we considered top 3 percentile of miRNA-gene interactions based on ProMISe scores. To run hiddenICP, we divided the BRCA patients into three random subsets of similar size to create the “different environments” required by the program. hiddenICP reports scores for all miRNA-gene pairs and we kept the top 2000 (top 3 percentile) of reported miRNA-gene interactions as the final interactions based on causal scores.

icpPAM50 also requires “different environments” based on the expression of PAM50 genes [40]. To address this requirement, we used the expression profiles of PAM50 genes from TCGA icpPAM50 to cluster the patients into multiple “environments”. Since icpPAM50 is not computationally efficient to run with high dimensional data, we used top 30 miRNAs and top 800 genes as input selected by the Feature Selection Based on The Most Variant Median Absolute Deviation (FSbyMAD) [41] method as used in [42]. We considered the top 2000 (top 8 percentile) of miRNA-gene interactions based on reported scores.

Since idaFast and jointIDA methods have high computational complexity and therefore are not scalable to large datasets, we run these two methods with top 50 miRNAs and top 1149 genes selected by using FSbyMAD. For idaFast, we selected top 2000 (top 3.5 percentile) of miRNA-gene interactions based on reported causal effect scores. We run jointIDA using R package *ParallelPC* [43]. In this case, we considered miRNA-gene interactions based on top 8.7 percentile of reported causal effect scores to get high confidence interactions.

We performed hypergeometric test to measure enrichment significance of computed miRNA targets in the known miRNA-gene interaction data (see Section 4.1). miRDriver had more significant miRNAs (*i.e*., hypergeometric test p-value ≤ 0.05) than all the other six methods (Table 2). These results indicate the performance of miRDriver in inferring accurate miRNA-gene interactions.

**Table 2:** Comparison results of miRDriver with six other methods. Eligible miRNAs are the ones that have at least one target in the ground truth dataset. Overlapping interactions and eligible miRNAs are computed with respect to miRDriver results.

| Method | miRDriver | AraCNe | hiddenICP | ProMise | idaFast | jointIDA | icpPAM50 |
| --- | --- | --- | --- | --- | --- | --- | --- |
| Input miRNAs | 255 | 255 | 255 | 255 | 50 | 50 | 30 |
| Input Genes | 6504 | 6504 | 6504 | 6504 | 1149 | 1149 | 800 |
| Selected miRNAs | 187 | 151 | 33 | 39 | 38 | 49 | 23 |
| Selected Genes | 776 | 266 | 6428 | 683 | 1052 | 1112 | 704 |
| Selected Interactions | 1430 | 278 | 2000 | 2000 | 2000 | 5000 | 2000 |
| Overlapping Interactions | N/A | 1 | 16 | 26 | 26 | 55 | 1 |
| Eligible miRNAs for Enrichment Test | 59 | 51 | 23 | 29 | 27 | 33 | 17 |
| Overlapping Eligible miRNAs | N/A | 35 | 7 | 25 | 21 | 27 | 16 |
| Method's Significant miRNAs in Overlap | N/A | 0 | 0 | 5 | 3 | 15 | 7 |
| miRDriver's Significant miRNAs in Overlap | N/A | 20 | 2 | 13 | 11 | 15 | 9 |

### 4.3 miRDriver produced significantly overlapping results on random subsets BRCA data

We tested the consistency of miRDriver results by splitting the BRCA data into two random patient groups. First, we miRDriver run on the entire patient cohort (*i.e., W*) then on two randomly chosen subsets, *S*_1_ and *S*_2_. We used the same input genes (*trans* genes) and input miRNAs (*cis* miRNAs) for all three sets. A few genes that had constant expression values among all the patients in a set were omitted from the LASSO step of miRDriver. We listed the number of input genes and miRNAs, and selected genes, miRNAs and miRNA-gene interactions for each dataset in Table 3. We found that miRDriver selected higher number of interactions and genes using set *S*_1_ than number of genes and interactions using set *W* and *S*_2_. Despite these differences, the overlap between them were significant. We performed the hypergeometric test to check the significance of selected overlapping miRNAs, selected genes and selected miRNA-gene interactions between each pair of these three sets and found all these overlaps were significant having hypergeometric p value ≤ 0.05 (Table 4). These results suggest that miRDriver could produce consistent results in different datasets of the same cancer type.

**Table 3:** miRDriver statistics when running it on the entire BRCA dataset (W) and two random subsets (*S*_1_ and *S*_2_).

| | Entire Set<br>($W$) | Subset<br>($S_1$) | Subset ($S_2$) |
| --- | --- | --- | --- |
| No. of input genes | 10,494 | 10,488 | 10,477 |
| No. of input miRNAs | 255 | 255 | 255 |
| No. of selected interactions | 1858 | 6551 | 1666 |
| No. of selected genes | 1114 | 2699 | 1059 |
| No. of selected miRNAs | 187 | 222 | 179 |

**Table 4:** Overlaps between selected interactions, genes and miRNAs when using miRDriver on the entire dataset (W), and random sub datasets (*S*_1_ and *S*_2_) in BRCA. All overlaps had significant hypergeometric p-value. The numbers in the parentheses show the overlap percentage with respect to *W*.

| Overlaps | $W \cap S_1$ | $W \cap S_2$ | $S_1 \cap S_2$ |
| --- | --- | --- | --- |
| Selected interactions | 756<br>(41%) | 560<br>(30%) | 454 |
| Selected genes | 603<br>(54%) | 461<br>(50%) | 526 |
| Selected miRNAs | 170<br>(91%) | 150<br>(80%) | 166 |

### 4.4 Selected miRNAs were enriched in the cancer-associated miRNAs

To test if the miRNAs selected by miRDriver were enriched in the cancer-related miRNAs, we used a list of 351 known oncogenic miRNAs from oncomiRDB (see Section 2.2.2). We performed Fisher’s exact test to check the association between the oncomiRDB miRNAs and selected miRNAs by miRDriver. For each cancer type, the background sets in the hypergeometric test consisted of all 1870 TCGA miRNAs among which 344 were common with oncomiRDB. In BRCA, 69 of 187 selected miRNAs and 275 of 1683 non-selected miRNAs were in oncomiRDB (p-value= 17.9e^−11^). In OV, 36 of 147 selected miRNAs were and 308 of 1723 non-selected miRNAs were in oncomiRDB (p-value = 0.03). The results indicate that miRDriver tends to select cancer-related miRNAs.

### 4.5 Selected miRNAs and genes were associated with survival of cancer patients

We performed multivariate survival analysis to assess the prognostic relevance of the selected and non-selected miRNAs by miRDriver. For breast cancer we considered age, race, HER2 status, estrogen status and progesterone status with the miRNA expression as independent variables and for OV age and race with miRNA expression were considered. In order to remove the confounding effect of other factors, we used the Adjusted Kaplan-Meier Estimator to compute adjusted survival curves by weighting the individual contributions by the inverse probability weighting (IPW) [44] using the R package *IPWsurvival* [45]. We considered two end points, Overall Survival (OS) and Progression Free Survival (PFI). In OS, patients who were dead from any cause considered as dead, otherwise censored. In PFI, patient having new tumor event whether it was a progression of disease, local recurrence, distant metastasis, new primary tumor event, or died with the cancer without new tumor event, including cases with a new tumor event whose type is N/A were considered as dead and all other patients were censored [35]. We computed prognostic miRNAs and genes while considering the entire patient cohort by dividing them into two groups as high expression and low expression of miRNA group. We found that a higher proportion of the selected miRNAs were prognostic miRNAs (log rank test p-value ≤ 0.05) than of the non-selected miRNAs (Table 5). To demonstrate the prognostic effect of the selected miRNAs in aberrated and non-aberrated patient groups, for each GISTIC region, we performed the survival analysis of the selected and non-selected miRNAs in that region within the frequently aberrated and non-frequently aberrated patient groups separately. We observed that selected miRNAs that were prognostic were higher in percentage in frequently aberrated patients than in non-frequently aberrated patients compared to non-selected miRNAs in BRCA for PFI end point and in OV for both OS and PFI end points except for when using OS as end point for BRCA (Table 6). Two examples of selected miRNAs from each cancer that have significant effect in aberrated patient group but not in the non-aberrated patient group are shown in Fig. 2. For instance, the patients in the frequently deleted group of the GISTIC region that harbors hsa-let-7g have worse survival when the hsa-let-7g is expressed low. The survival curves for all prognostic selected miRNAs in aberrated and non-aberrated groups are in the Supplementary File 1.

**Table 5:** Percentage of prognostic selected miRNAs and prognostic non-selected miRNAs.

| End | Point | Prognostic miRNAs in<br>selected miRNAs | Prognostic miRNAs in<br>non-selected miRNAs |
| --- | --- | --- | --- |
| OS in BRCA |  | 18% | 13% |
| PFI in BRCA |  | 9.6% | 4.4% |
| OS in OV |  | 10% | 8.6% |
| PFI in OV |  | 11% | 6% |

**Table 6:** Percentage of prognostic selected miRNAs vs. prognostic non-selected miRNAs within the frequently aberrated and not-frequently aberrated patient groups.

| End | Point | Significant selected<br>miRNAs in aberrated<br>but not in non-aberrated | Significant non-selected miRNAs in<br>aberrated but not in<br>non-aberrated |
| --- | --- | --- | --- |
| OS in BRCA |  | 8.6% | 10% |
| PFI in BRCA |  | 4.3% | 2.9% |
| OS in OV |  | 7.2% | 6.8% |
| PFI in OV |  | 7.2% | 3.7% |

**Figure 2:**
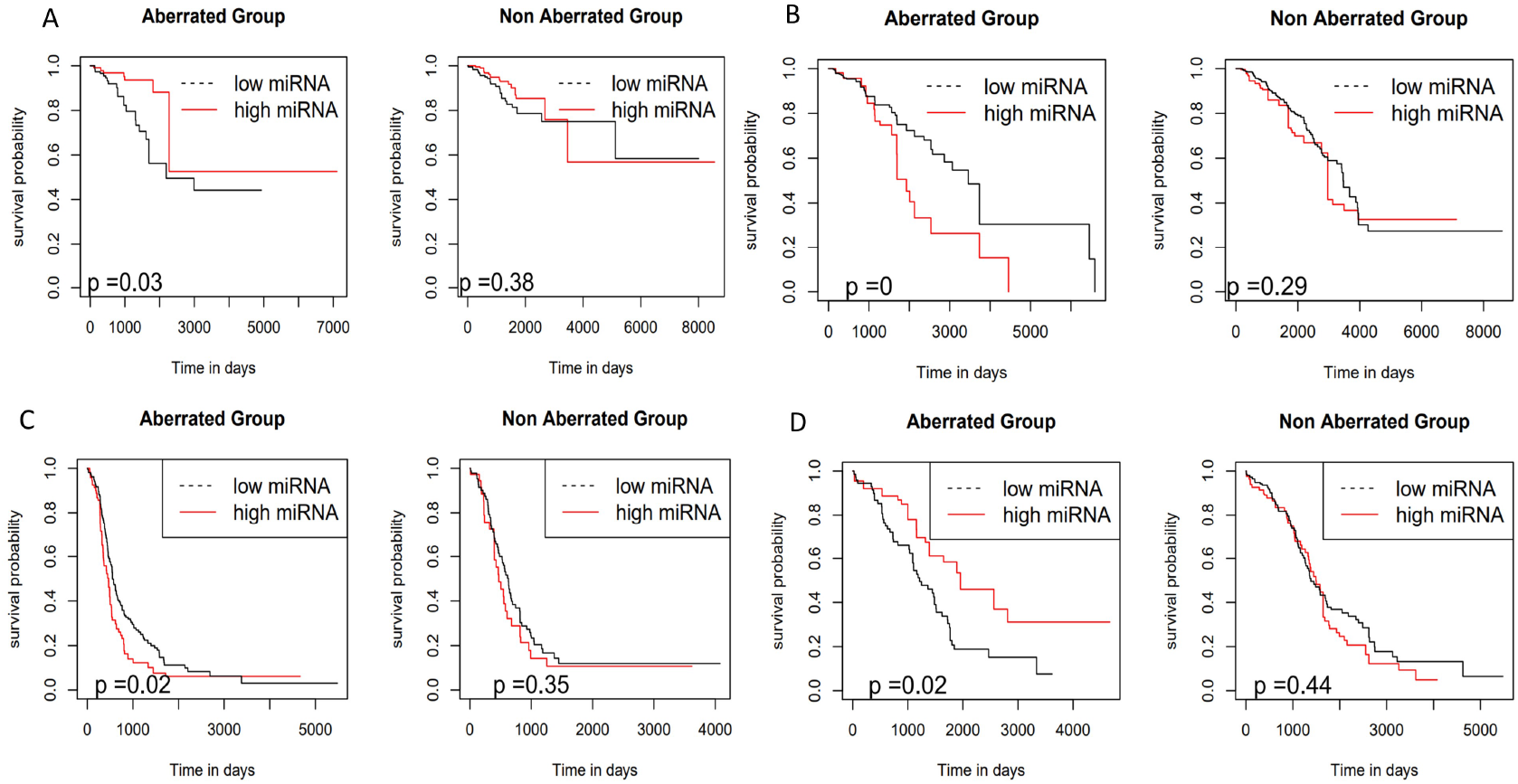
Adjusted Kaplan-Meier plots for selected miRNAs in aberrated and non-aberrated patient groups. A) hsa-let-7g in a deleted region in BRCA. B) hsa-mir-10a in an amplified region in BRCA C) hsa-mir-3193 in an amplified region in OV D) hsa-mir-443 in a deleted region in OV. In each case, miRNAs are significant prognostic factors only within frequently aberrated patient group.

We computed the hazard ratio (HR) of higher degree selected genes (genes targeted by four or more miRNAs) and lower degree selected genes (genes targeted by three or fewer miRNAs) using multivariate Cox regression analysis [46]. The genes with Cox regression p-value ≤ 0.05 were considered as prognostic genes. A ln (HR) > 0 implies that an increase of expression of the gene increases the risk of an event, while a ln (HR) < 0 implies that an increase of the gene expression decreases the risk of an event.

We compared the median of the ln (HR) of each group using the Wilcoxon rank-sum test. In BRCA with OS, interestingly higher degree genes possessed lower hazard ratios whereas in OV with PFI, higher degree genes possessed higher hazard ratios compared to lower degree genes (Fig. 3A, B).

**Figure 3:**
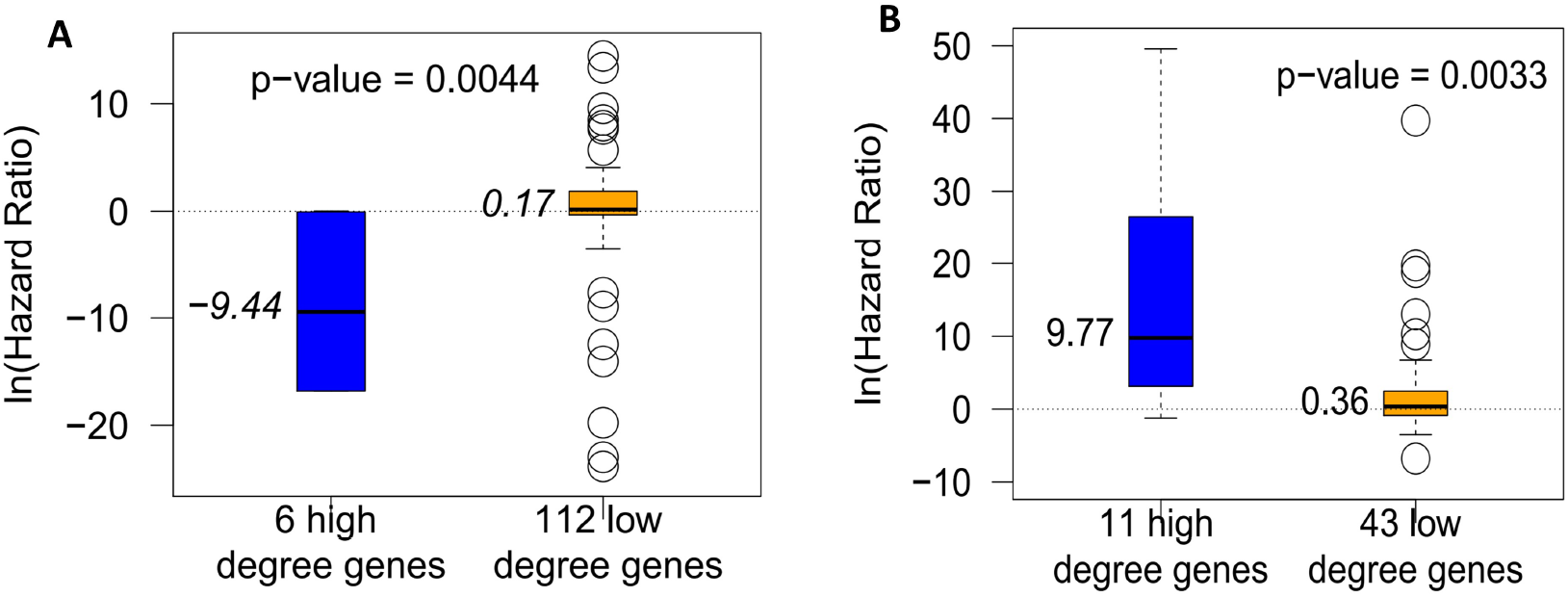
Plots of natural logarithm of hazard ratios in high degree and low degree genes, (A) Breast cancer with PFI, (B) Ovarian cancer with OS.

## 5 Discussion

We developed a computational pipeline called miRDriver, which integrates multi omics datasets such as copy number variation, DNA methylation, transcription factors, gene and miRNA expression to infer copy number-derived miRNA-gene interactions in cancer. We applied miRDriver on breast and ovarian cancer datasets from TCGA. In each case, miRDriver was able to select many miRNA-gene interactions that were enriched in known miRNA-target databases. We observed that selected miRNAs by miRDriver were significantly enriched in the known cancer-related miRNAs. The selected miRNAs that targeted many genes, for instance miR-23a, miR-27b, miR-181d in ovarian, miR-1224, miR-31, let-7g in breast cancer and let-7b in both cancers are known to be involved in cancer [48–51]. Hu et al. found that miR-23a has role in promoting tumor growth and apoptosis via target gene PDCD4 in gastric cancer [54]. A recent study found association of miR-1224 in tumor metastasis in gastric cancer [57]. It was found that let-7 family is related with tumorigenesis in breast cancer [55]. Studies on deregulated let-7 expression and its role in tumorigenesis have been conducted [56].

We evaluated the prognostic value of miRDriver’s selected miRNAs vs non-selected miRNA’s using Adjusted Kaplan-Myer survival analysis. We found that percentage of prognostic miRNAs was greater in selected miRNAs than in non-selected miRNAs, which indicates that miRDriver tends to select prognostic miRNAs. Also, percentage of prognostic miRNAs was higher in copy number aberrated patient cohort than in non-aberrated regions, which suggests that copy number aberration event tends to increase the prognostic value of miRNAs.

In BRCA, the selected high degree genes (i.e., genes targeted by ≥4 miRNAs) had low hazard ratios (HR) (median ln (HR) = −9.44) in patient’s survival with progression free interval (see Section 4.5 & Fig. 3A). This result suggests that these genes tend to be acting as tumor suppressor genes. Among these high degree genes, CLIC6 was found to be associated in ion channels, which are implicated in breast cancer [58]. In a recent work [59], the mechanism of CLIC proteins in promoting an aggressive invasive carcinoma was explored. Another selected high degree gene in BRCA, PALM2 showed differential expression in chemotherapy response in patients with metastatic colorectal carcinoma [60]. Conversely, in OV, the selected high degree genes had high hazard ratios (HR) (median ln (HR) = 9.77) in overall survival of cancer patients (Fig. 3B). This result suggests that these high degree genes tend to be acting as oncogenes. In OV, high degree genes like MEP1B was reported to be associated with proliferation and invasion of colorectal cancer [61] and LINC00189 reported to be a possible biomarker of the bladder cancer [62]. These studies support the idea of exploring these high degree genes as tumor suppressor genes and oncogenes. These high degree genes can serve as potential biomarkers to predict prognosis in cancer patients.

We compared miRDriver’s LASSO steps with network-based methods AraCNe and ProMIse, and causal inference-based methods such as hiddenICP, ICP-PAM50, idaFast and jointIDA. We showed that miRDriver substantially outperformed other tools in selection of significantly known miRNA-gene interactions (Table 2). We also checked the consistency of miRDriver results in different subsets of breast cancer patients and found significant overlaps between different results (Table 3).

In miRDriver, we integrated high dimensional multi omics data and it possessed some computational challenges. To control this, we ran our LASSO step in parallel clusters. For breast cancer, we ran 1,049,400 LASSO regressions using 210 cores Dual Intel Processors with 512 GB RAM. Despite of that, due to large number of patients and *trans* genes in breast cancer data, it took five days to complete the computation. Whereas, for ovarian cancer, the computation was finished within 2 hours due to fewer patients and *trans* genes.

For the future versions of miRDriver, more datasets such as mutations, histone modification and chromatin accessibility datasets such as ATAC-seq could be incorporated. The current version of miRDriver does not incorporate the competing endogenous RNA (ceRNA) interactions [64]. These interactions could affect how miRNAs interact with their target genes. When computing potential targets of miRNAs, putative ceRNA interactions could be computed simultaneously.

In conclusion, we presented miRDriver, a computational method that integrates genomic, transcriptomic, and epigenetic datasets to infer copy number derived miRNAs that affect downstream target genes in cancer. This could help find genes which could be therapeutic targets of drugs and opens the opportunity to establish novel biomarkers.

