## Supplemental File 1 for "miRDriver: A Tool to Infer Copy Number Derived miRNA-Gene Networks in Cancer"

### ADJUSTED KAPLAN-MEIER SURVIVAL CURVES IN BRCA

#### Amplification Peak 12

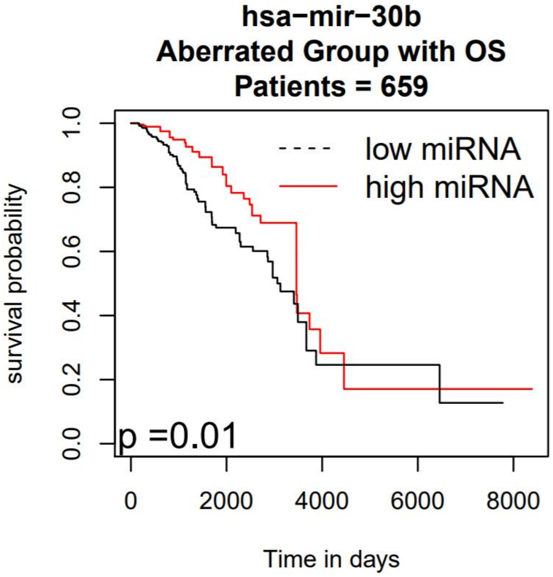


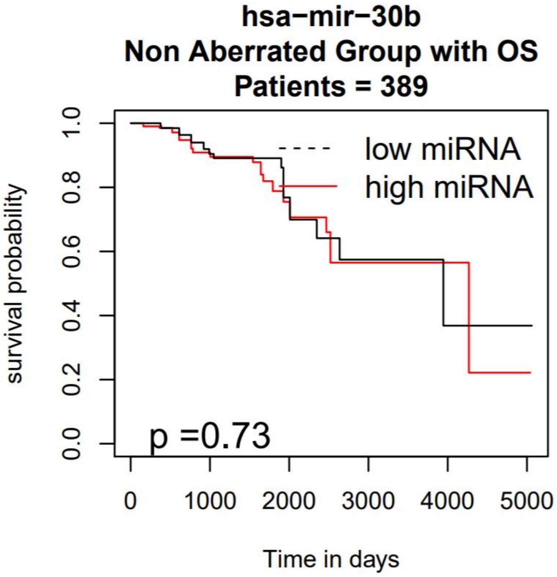


#### Amplification Peak 12

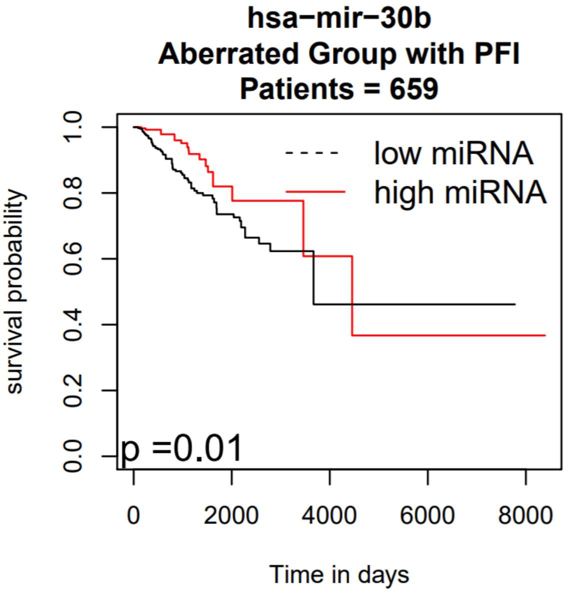


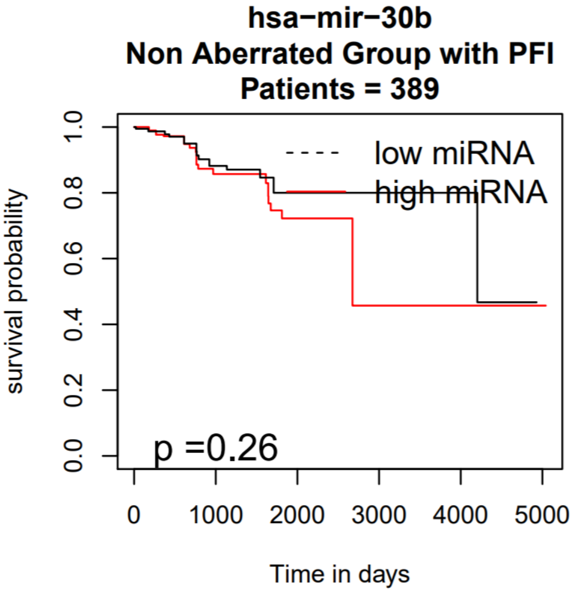

### Amplification Peak 12

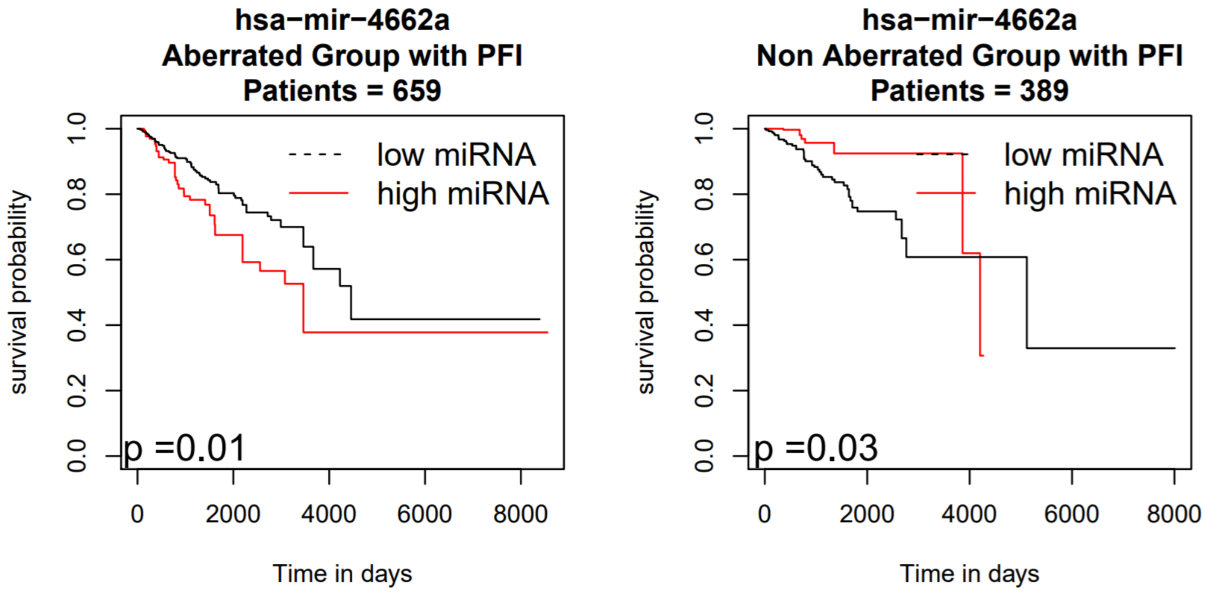


### Amplification Peak 12

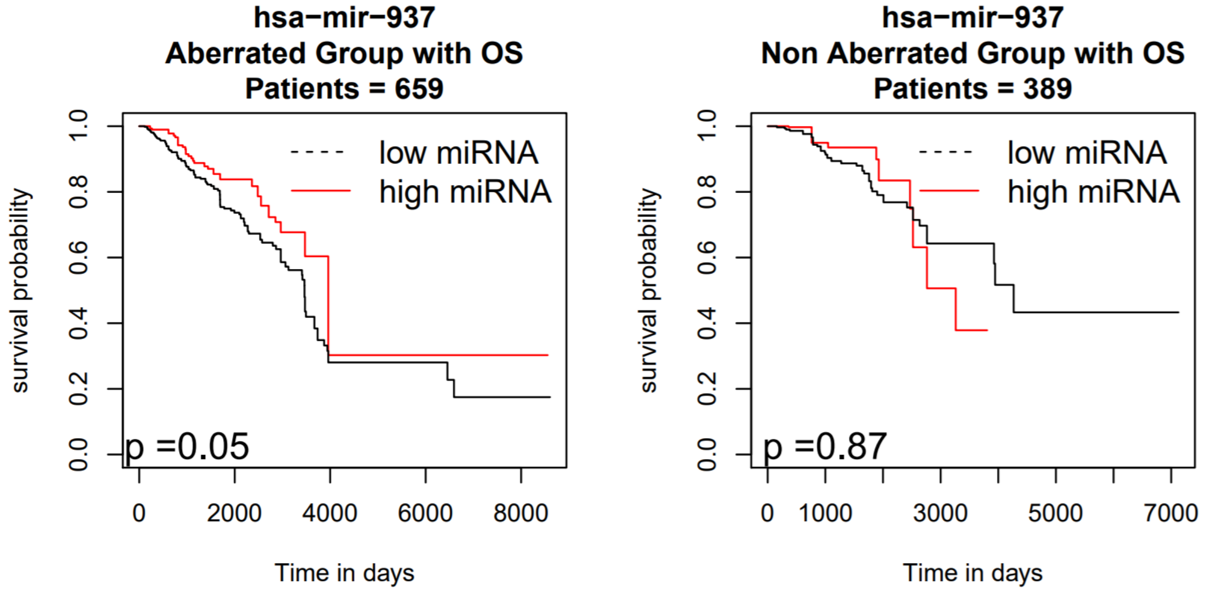

### Amplification Peak 24

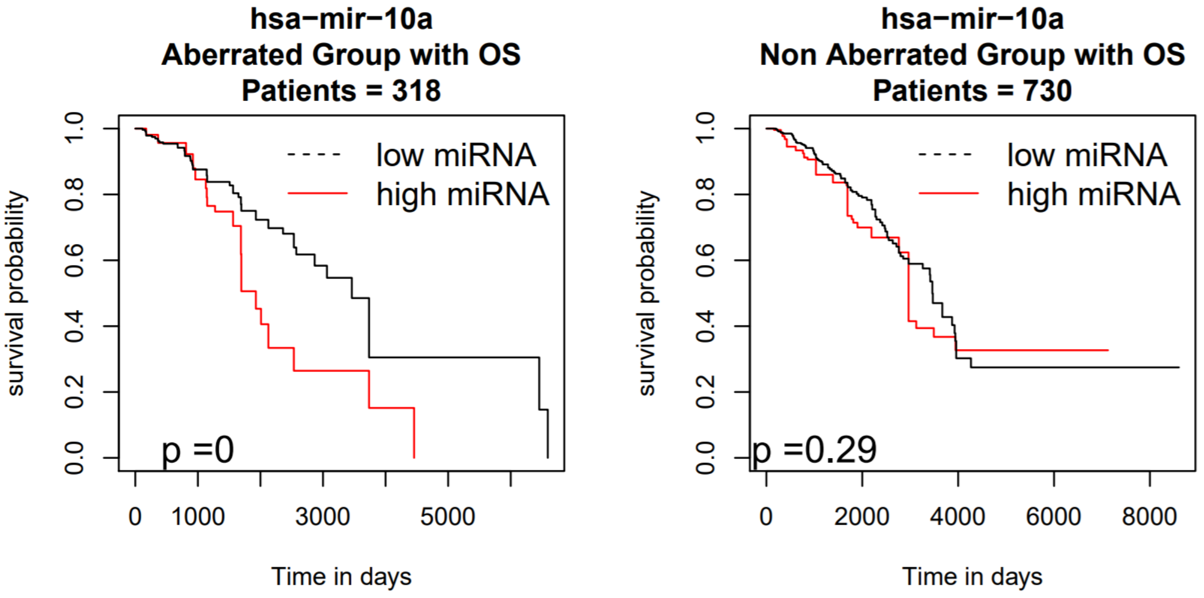


### Amplification Peak 24

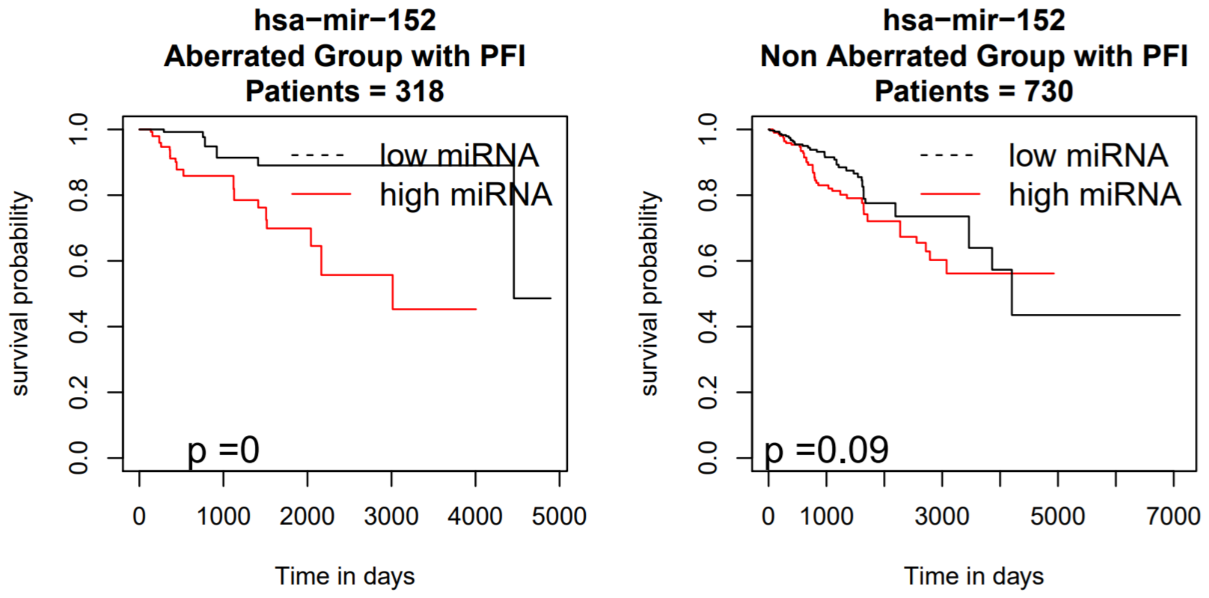

### Amplification Peak 24

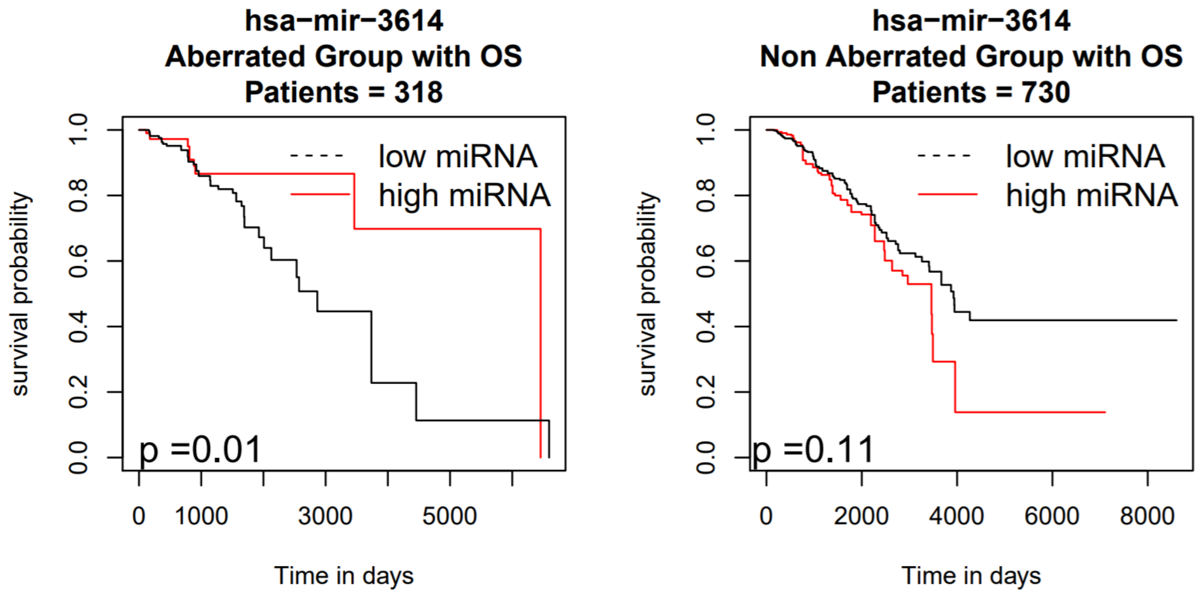


### Amplification Peak 29

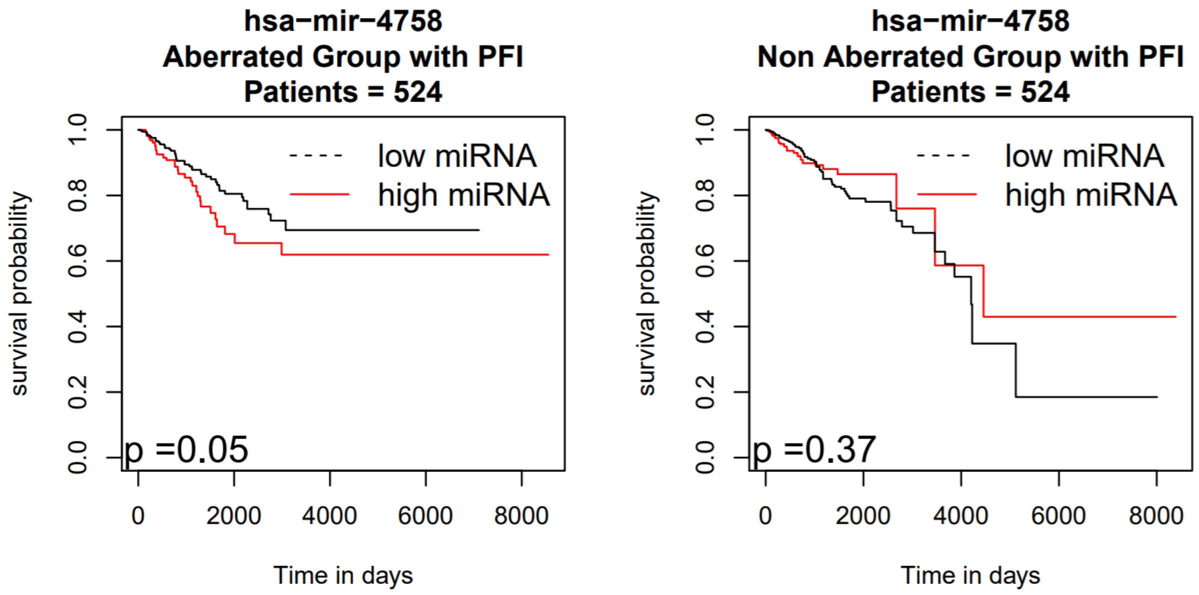

### Deletion Peak 7

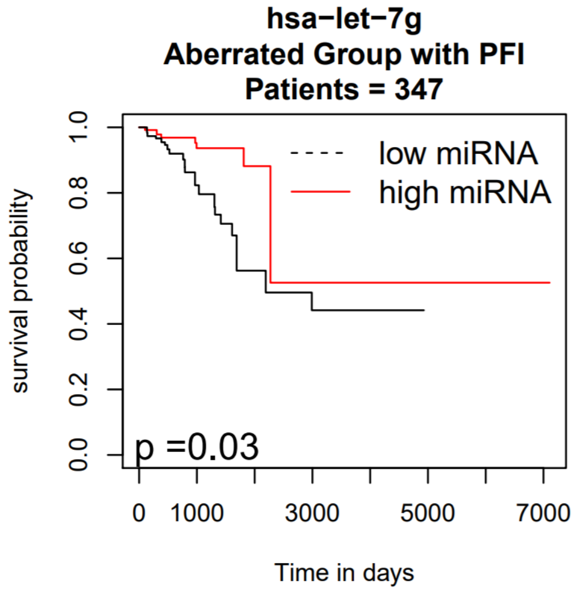


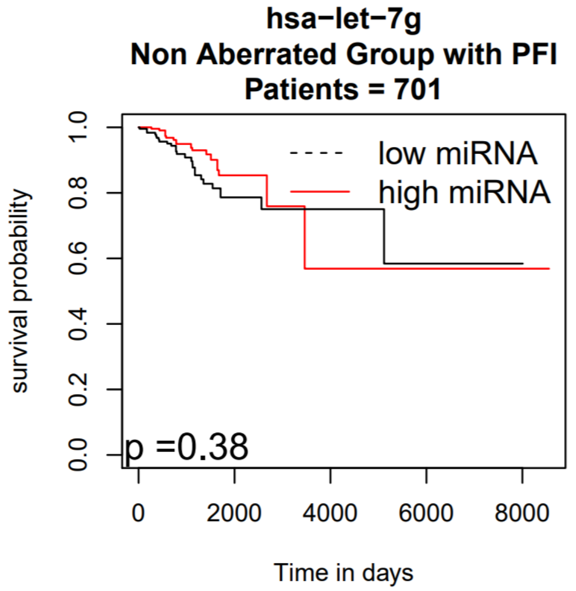


### Deletion Peak 7

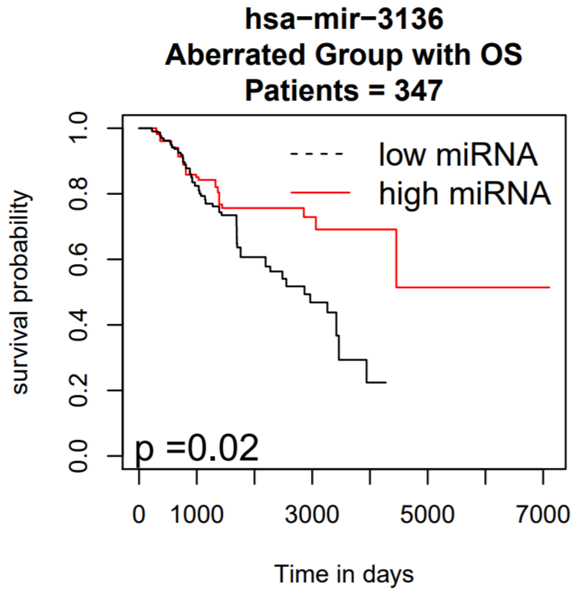


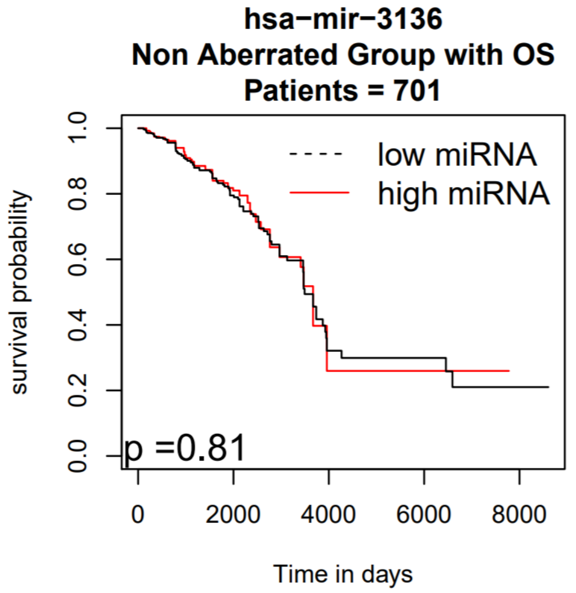

**Deletion Peak 17**

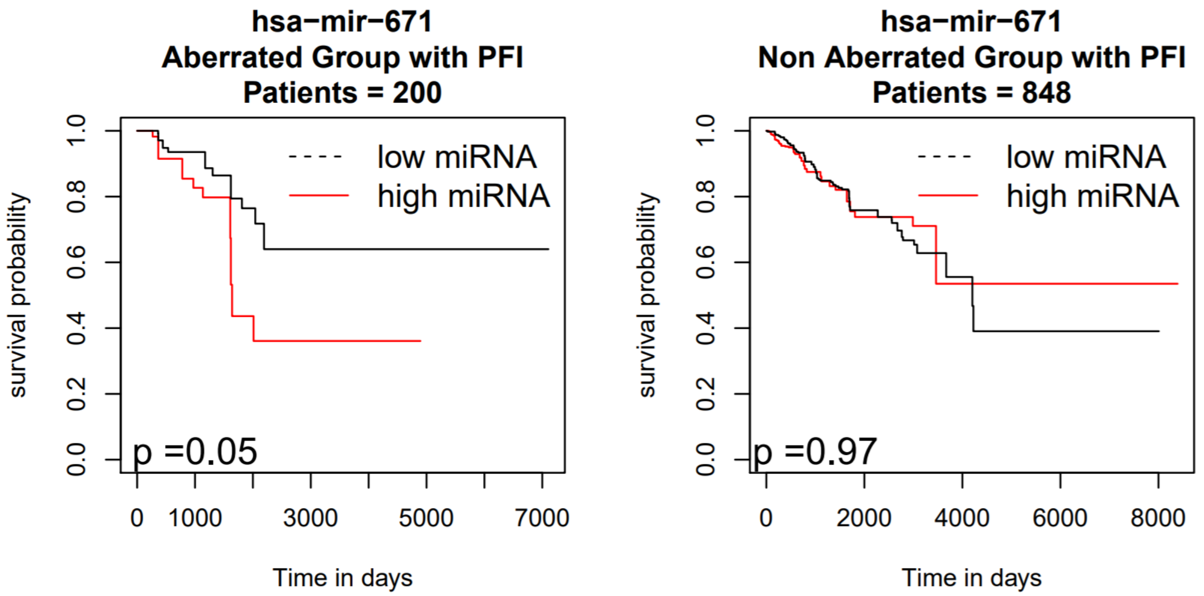


**Deletion Peak 24**

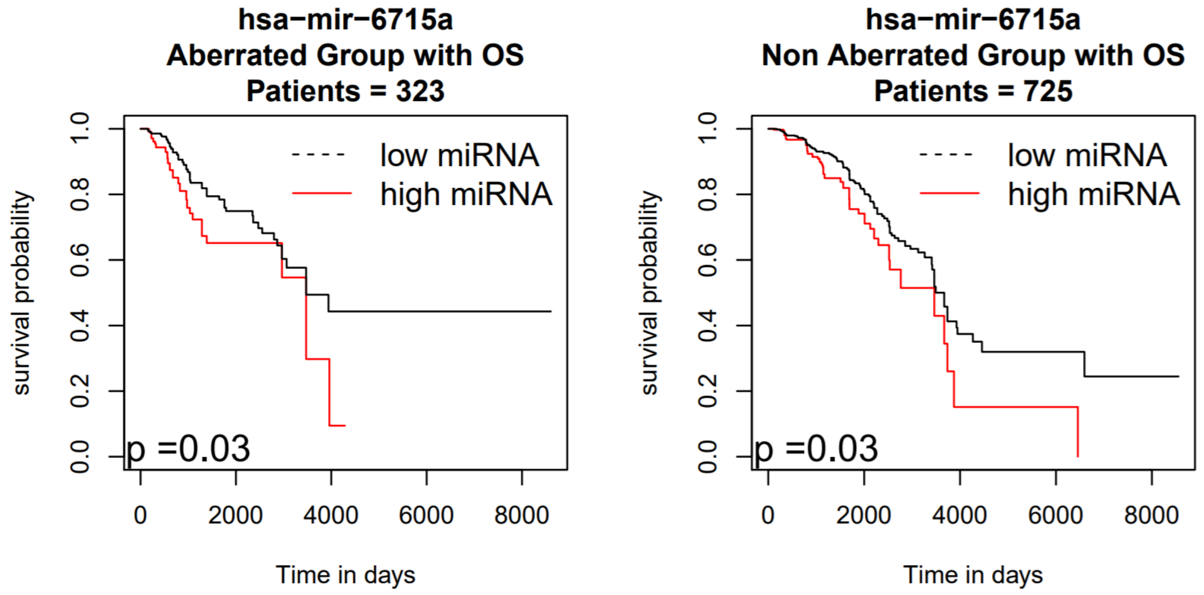

### Deletion Peak 27

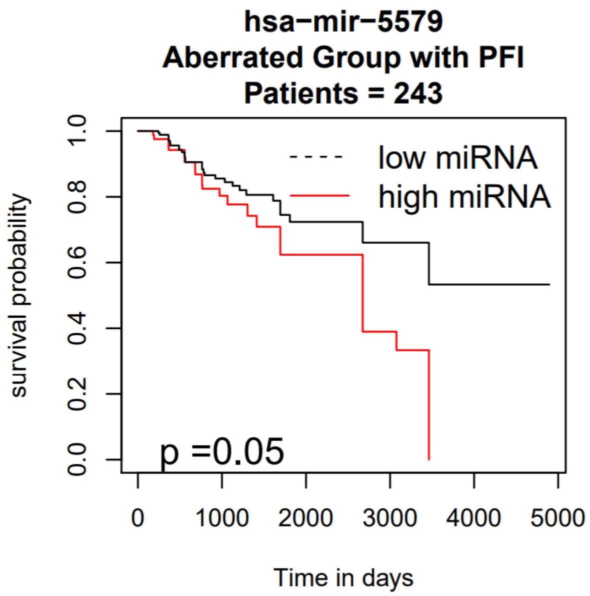


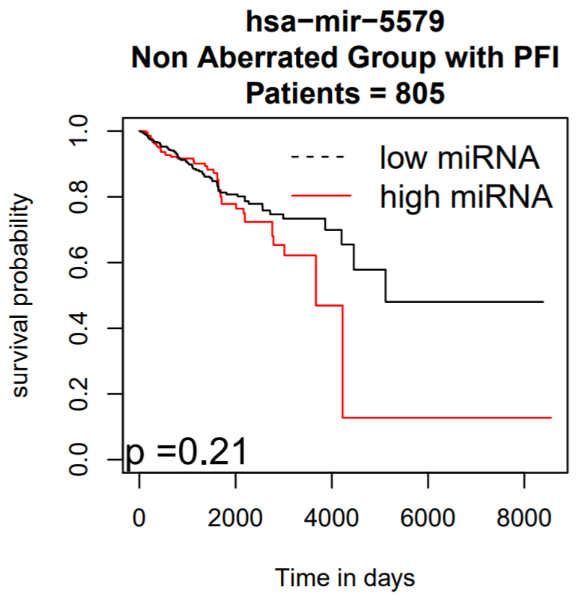


### Deletion Peak 28

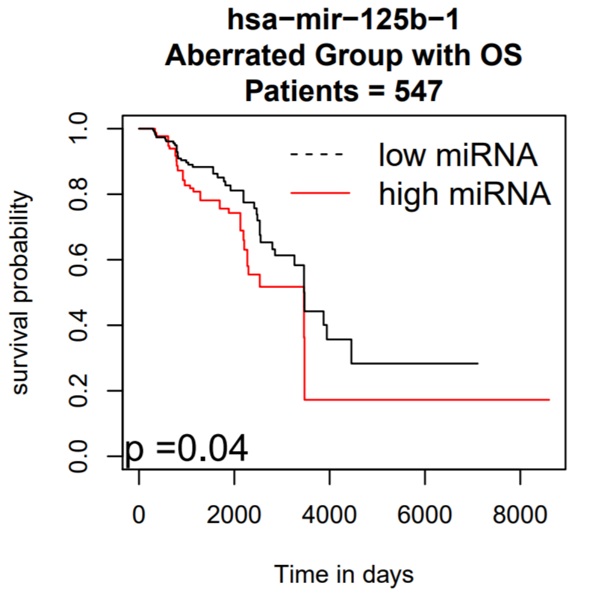


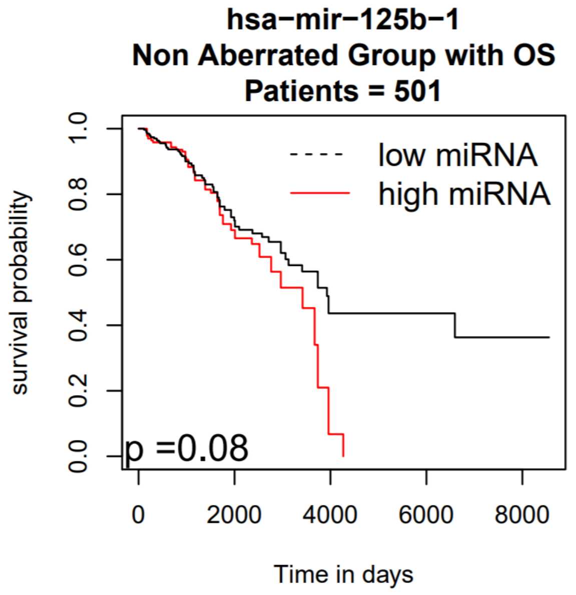

### Deletion Peak 28

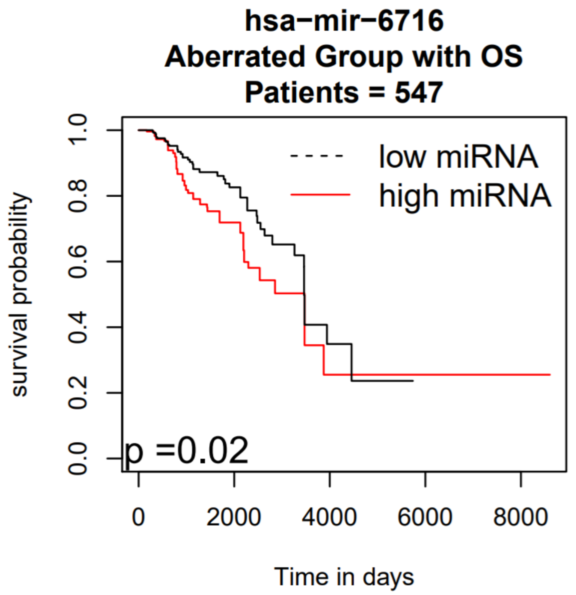


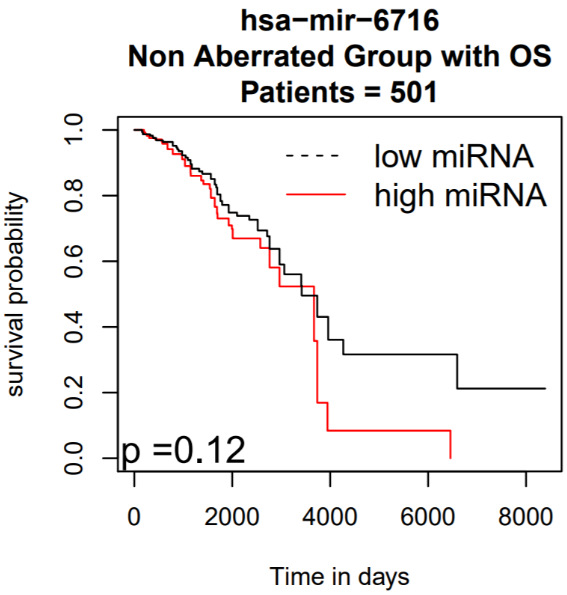


### Deletion Peak 36

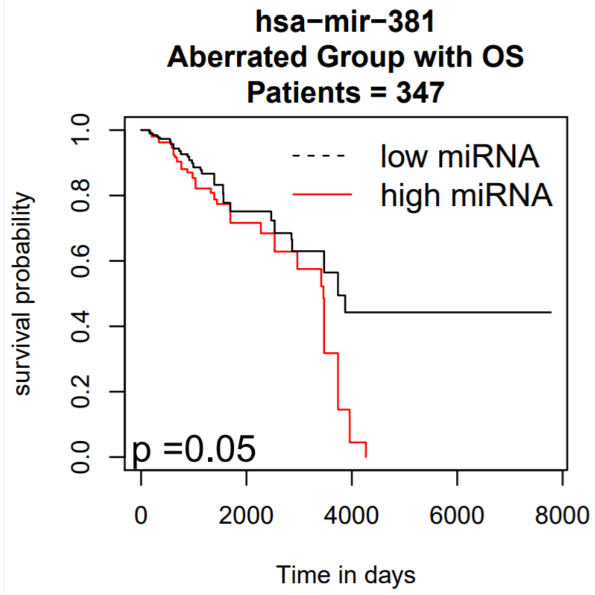


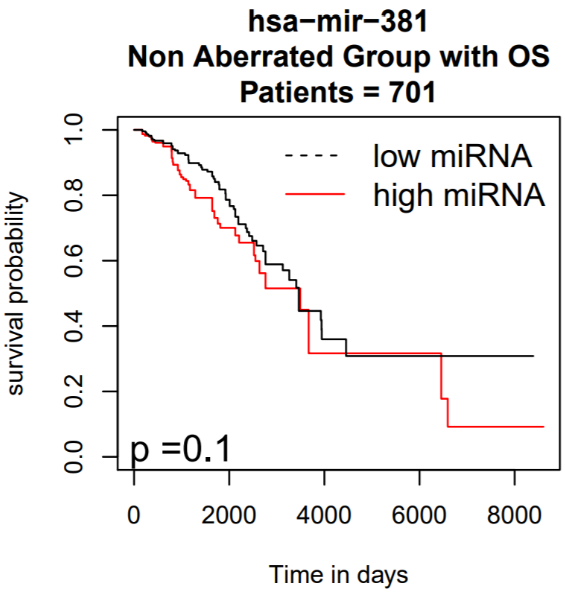

**Deletion Peak 36**

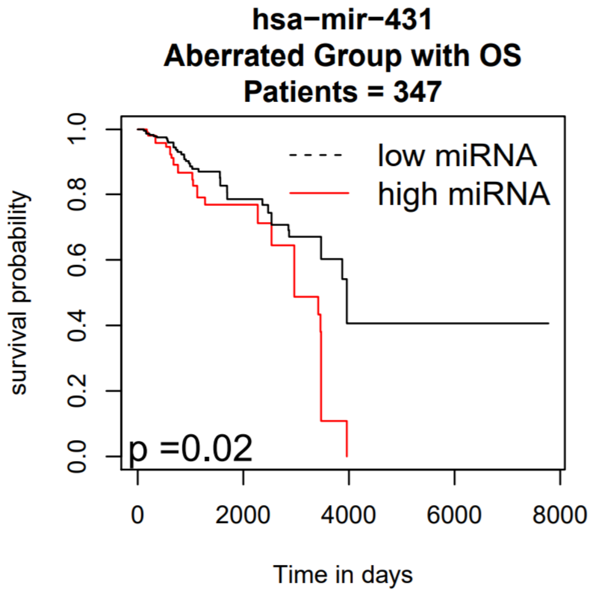


**Deletion Peak 36**

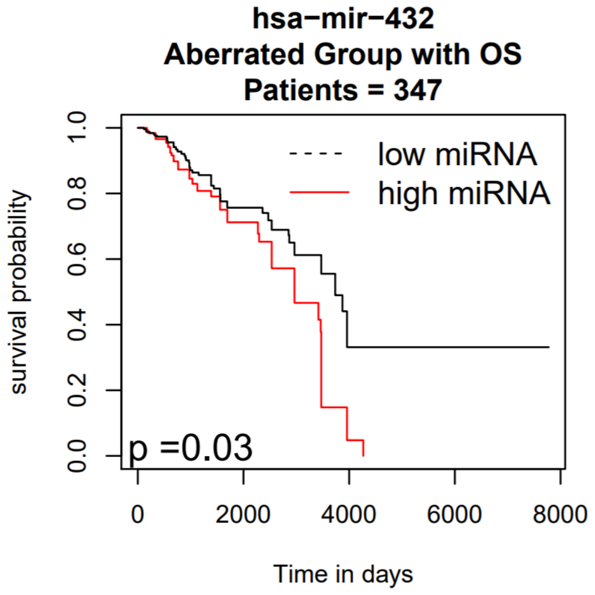

### Deletion Peak 36

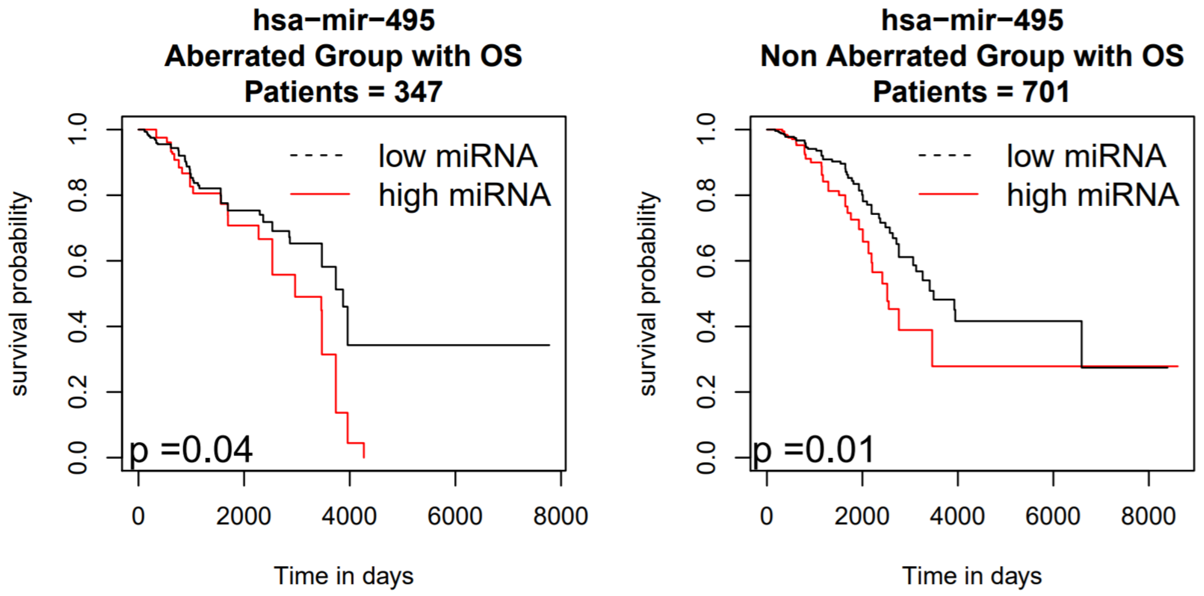


### Deletion Peak 38

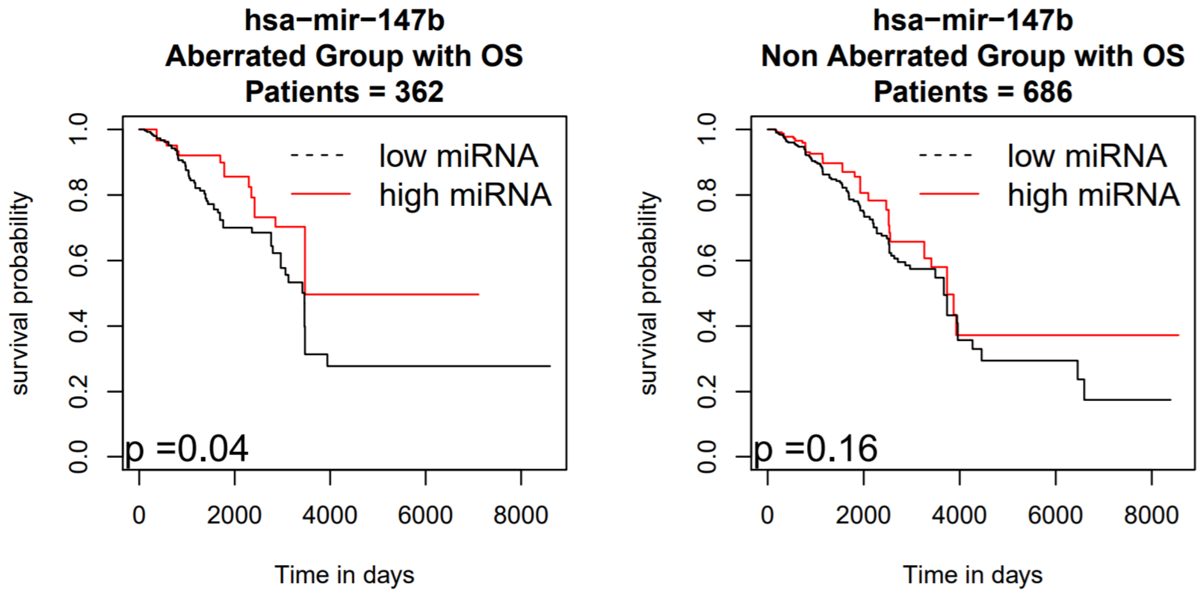

### Deletion Peak 38

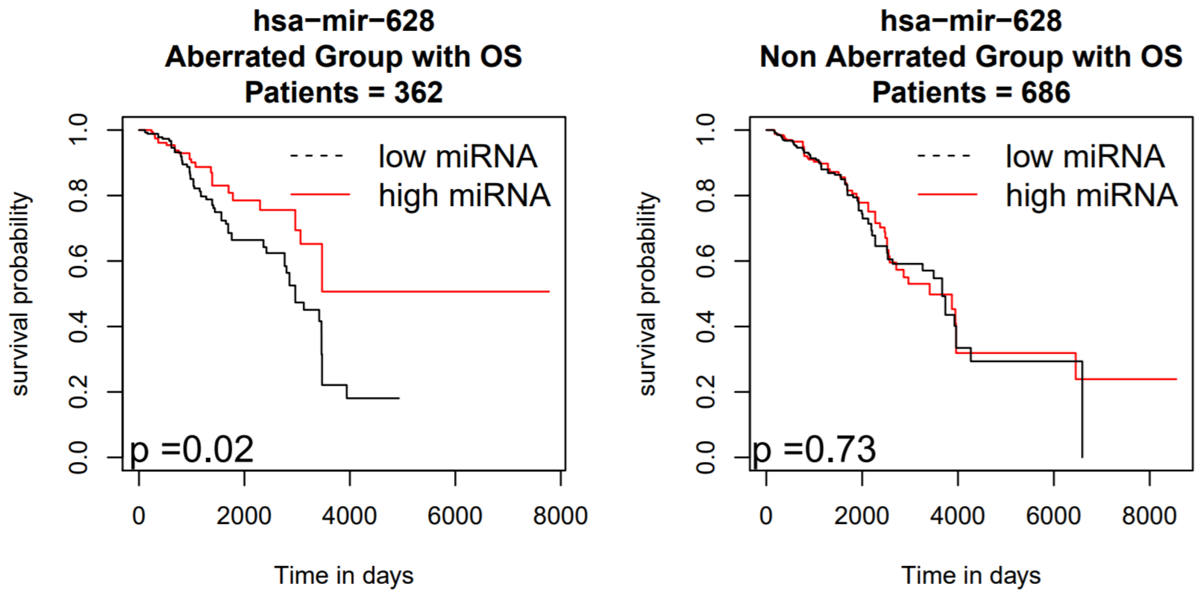


### Deletion Peak 53

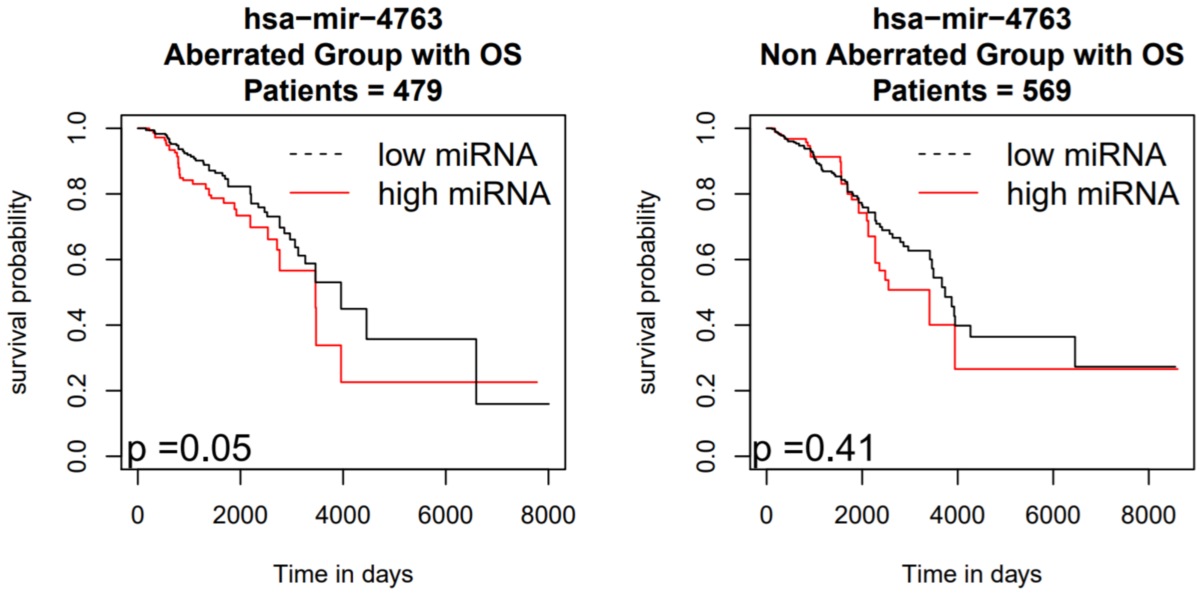

### ADJUSTED KAPLAN-MEIER SURVIVAL CURVES IN OV

#### Amplification Peak 25

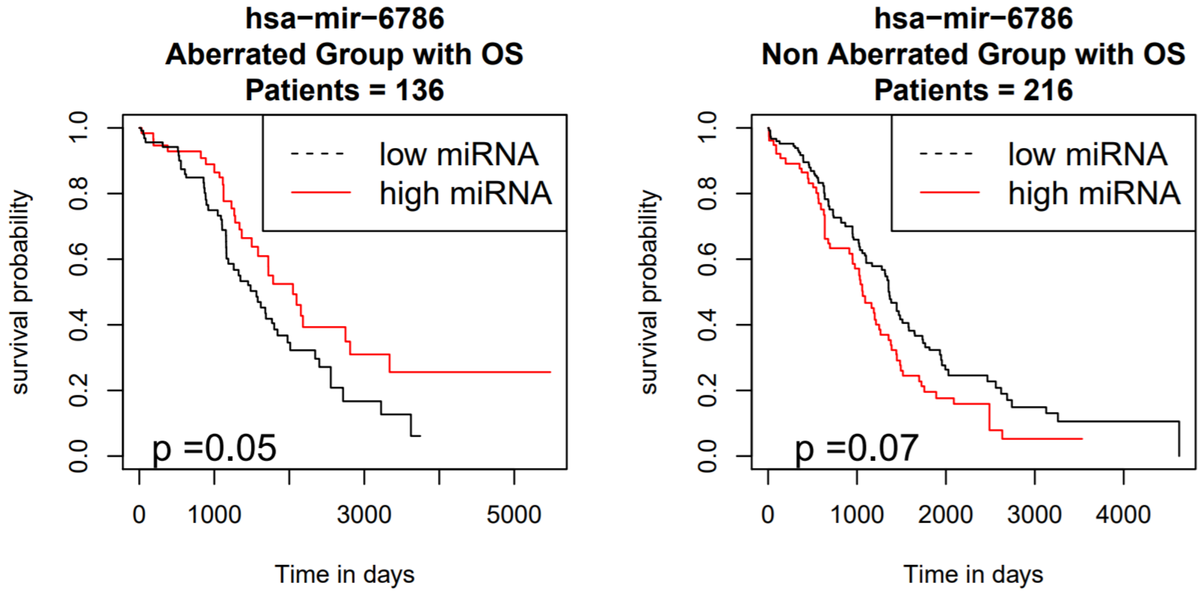


#### Amplification Peak 32

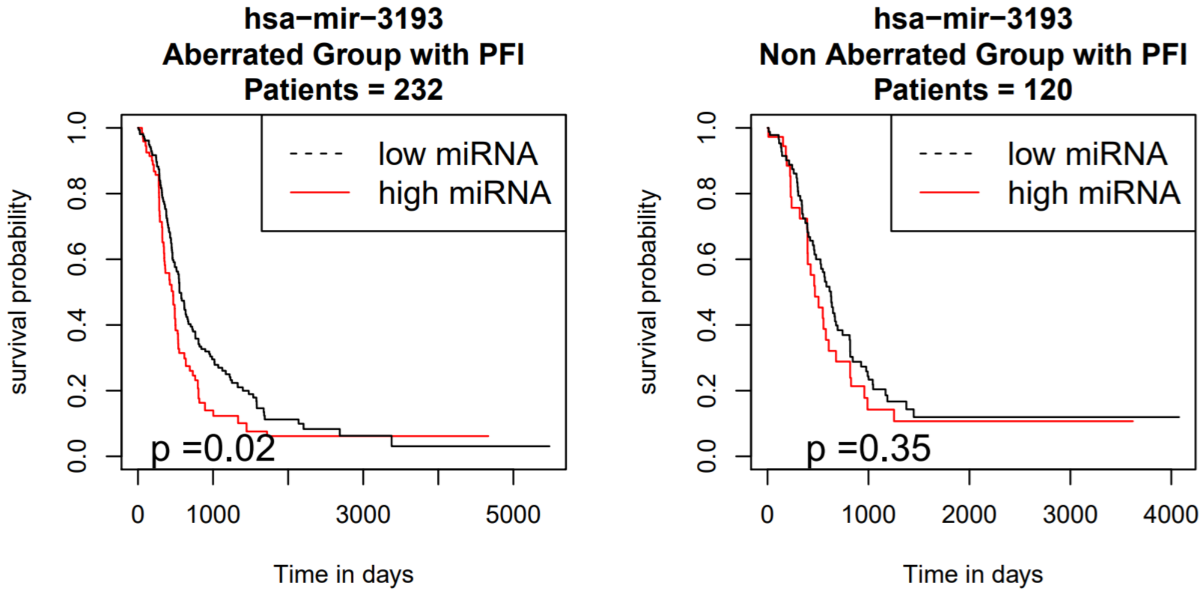

### Amplification Peak 33

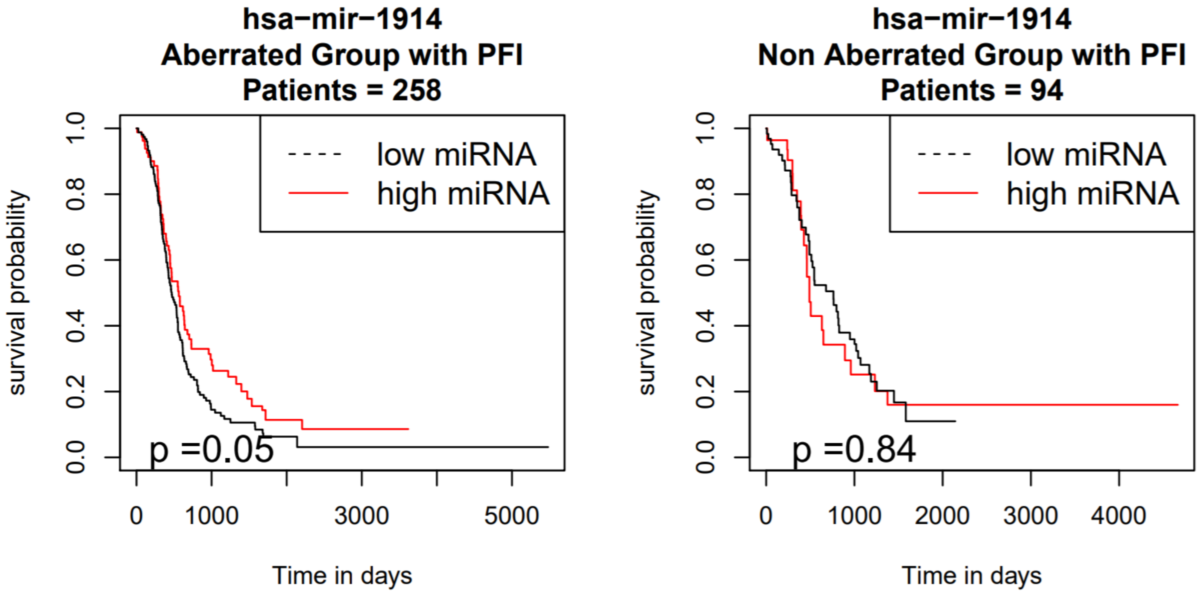


### Amplification Peak 33

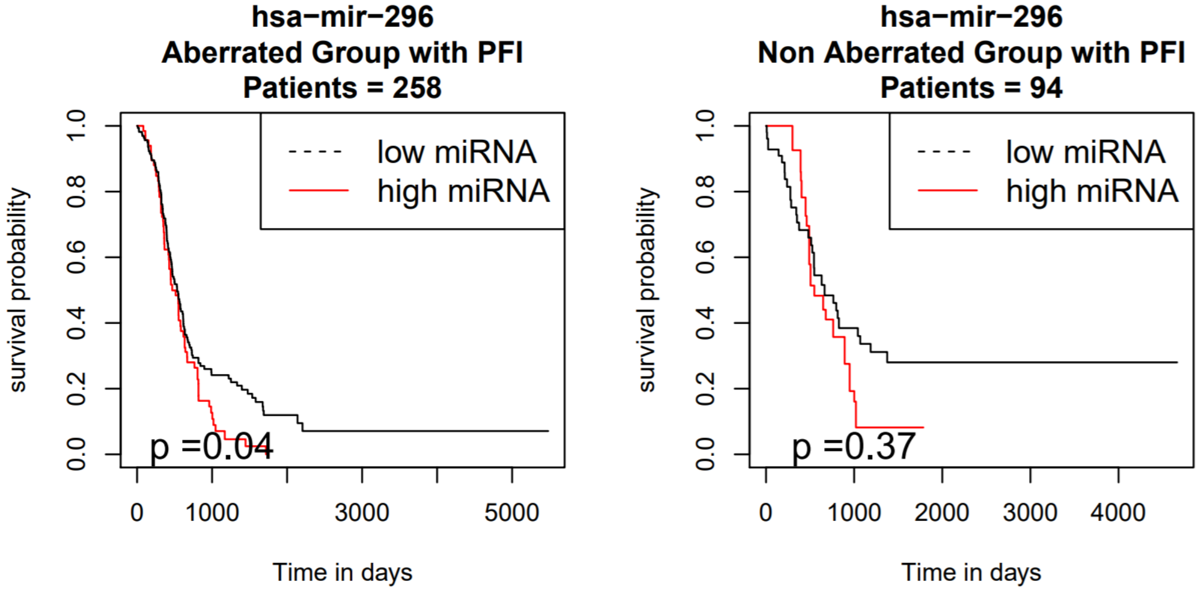

### Amplification Peak 33

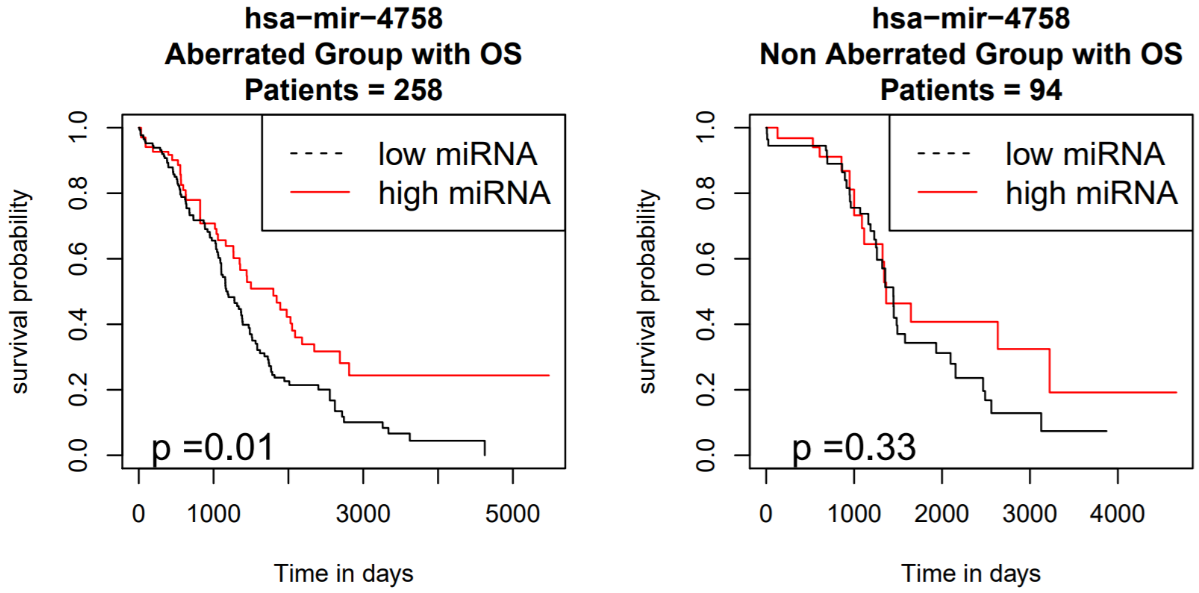


### Amplification Peak 33

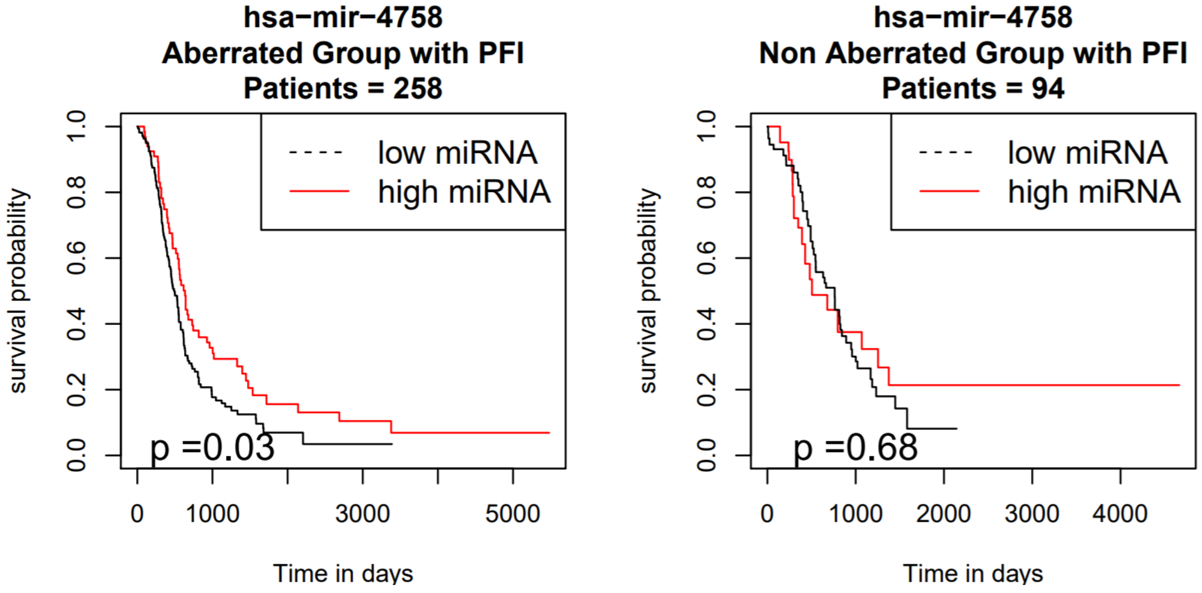

**Deletion Peak 5**

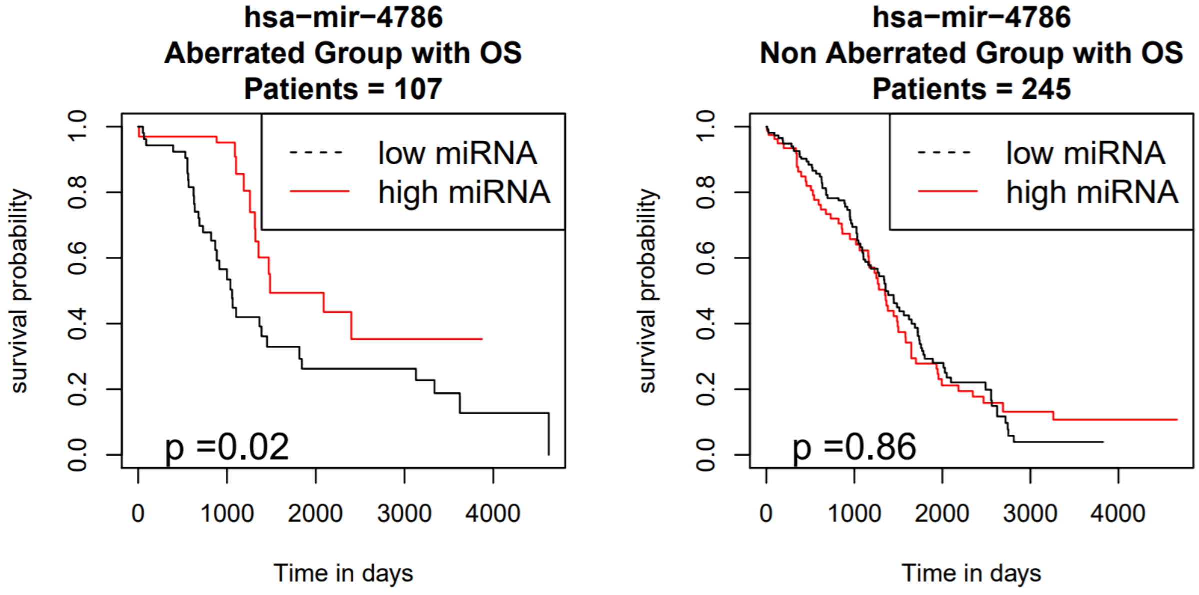


**Deletion Peak 5**

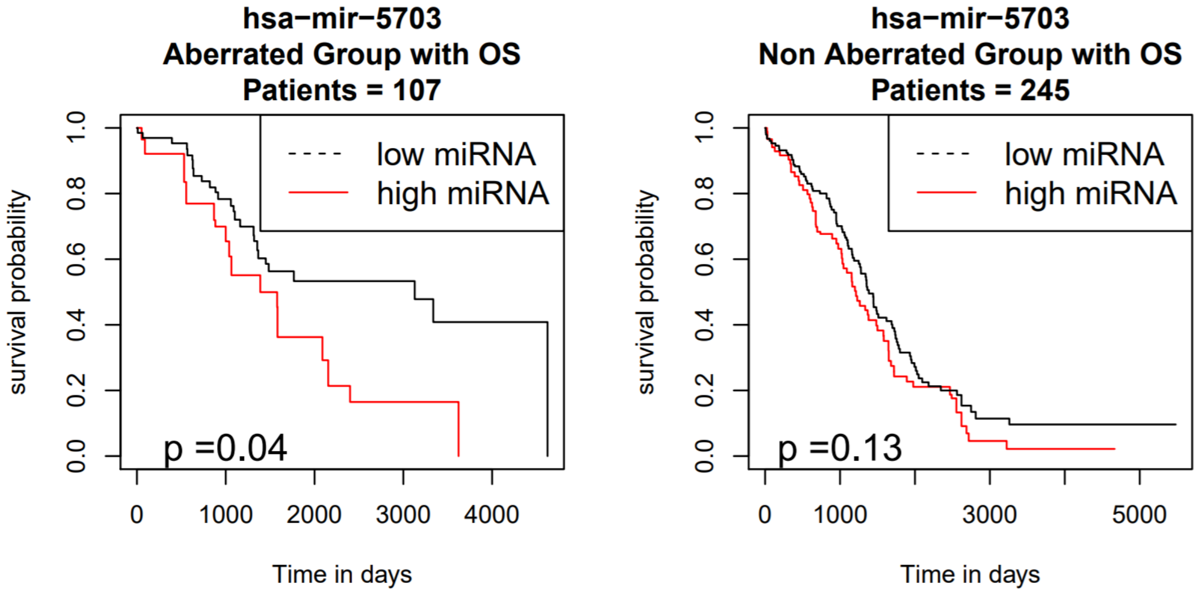

### Deletion Peak 6

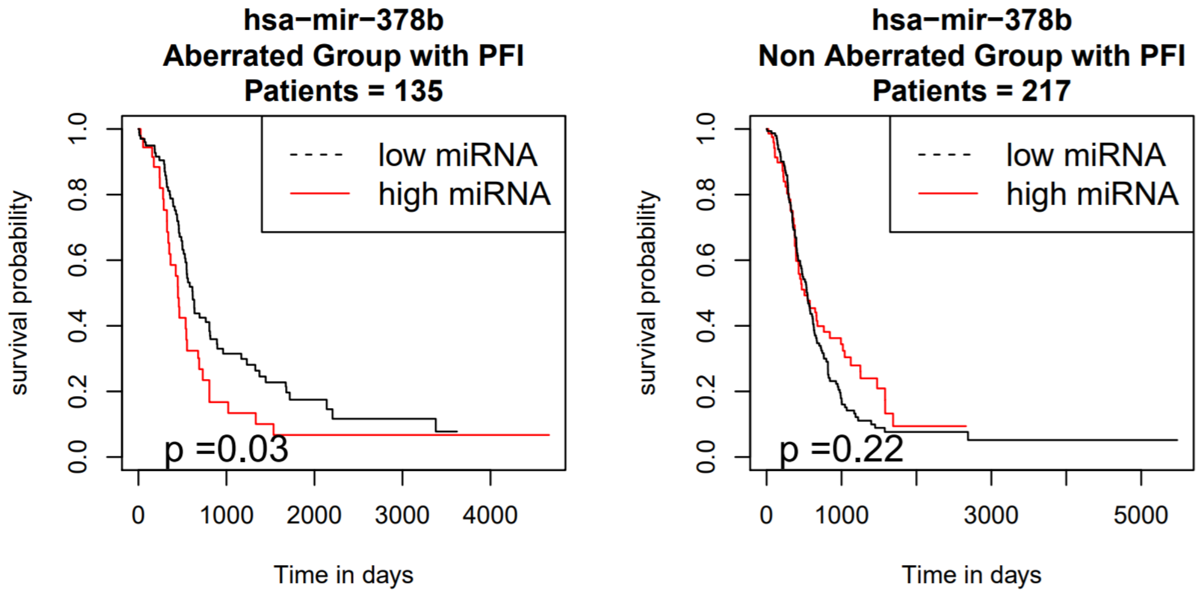


### Deletion Peak 7

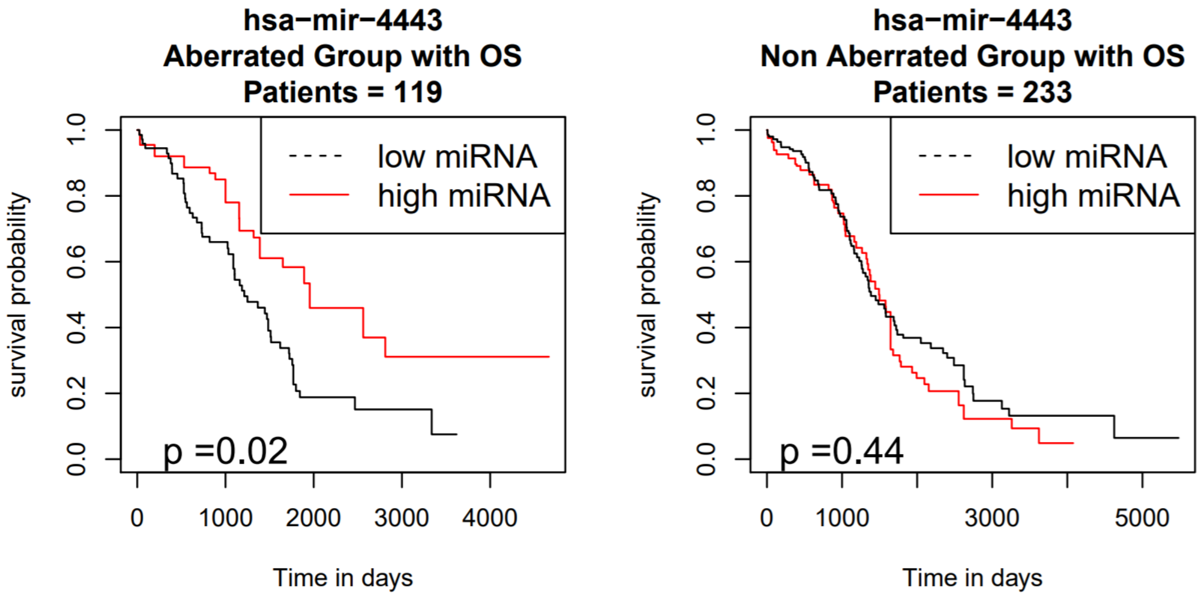

### Deletion Peak 7

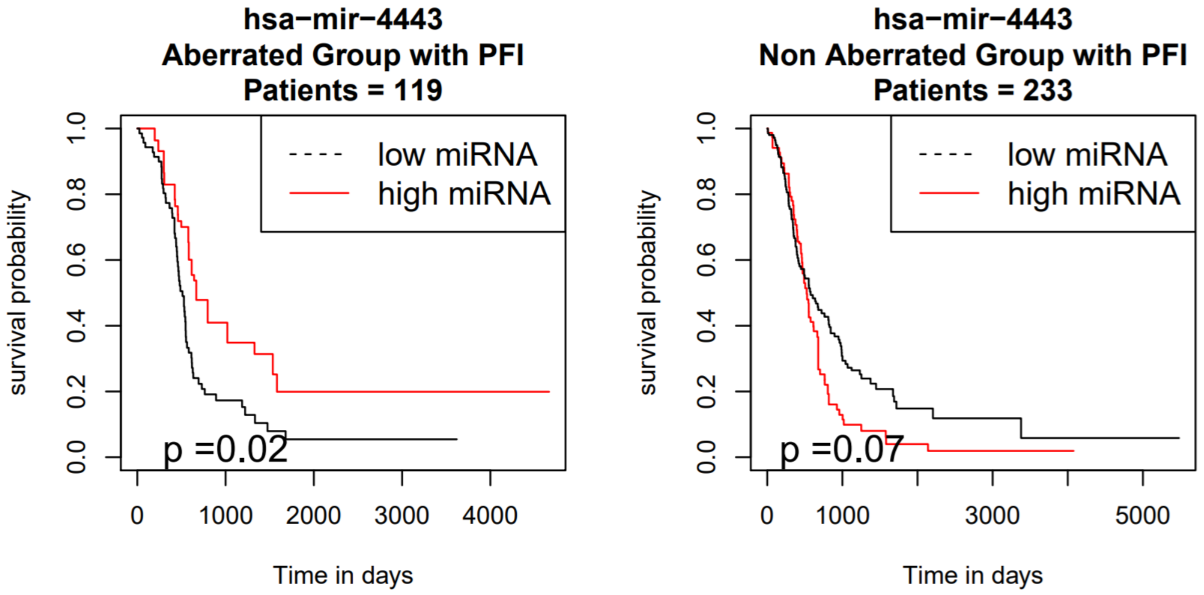


### Deletion Peak 7

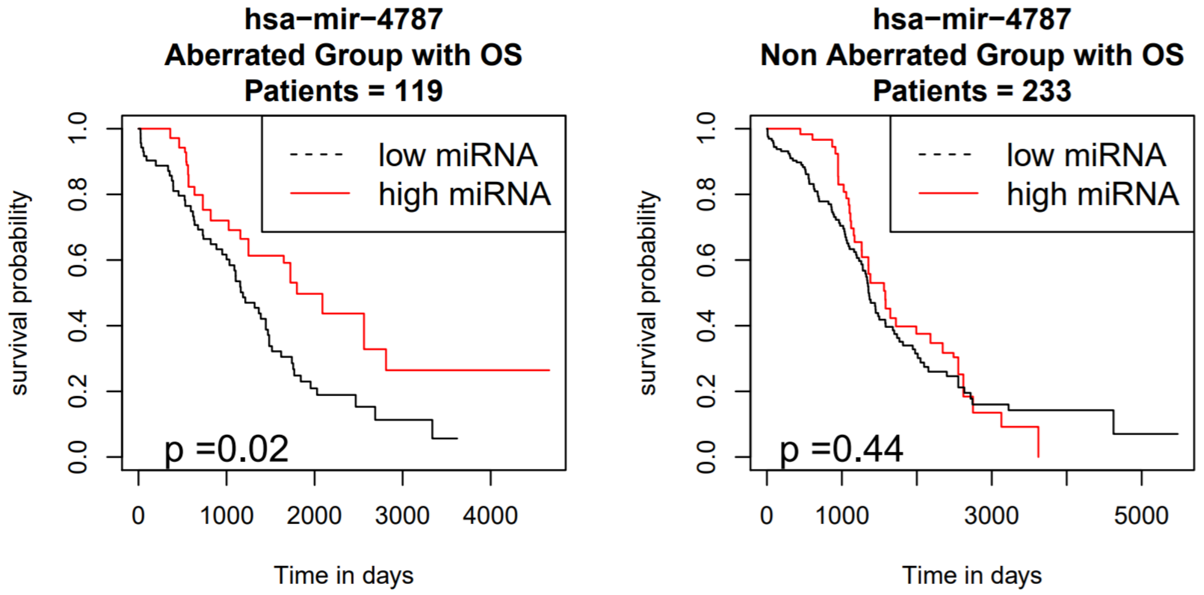

### Deletion Peak 7

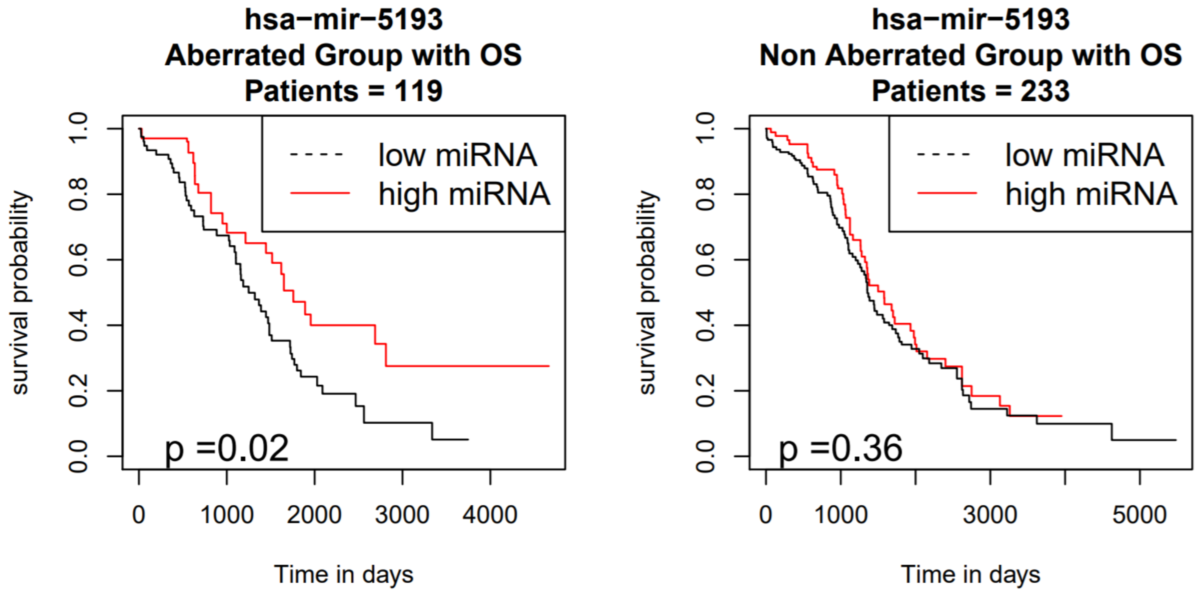


### Deletion Peak 7

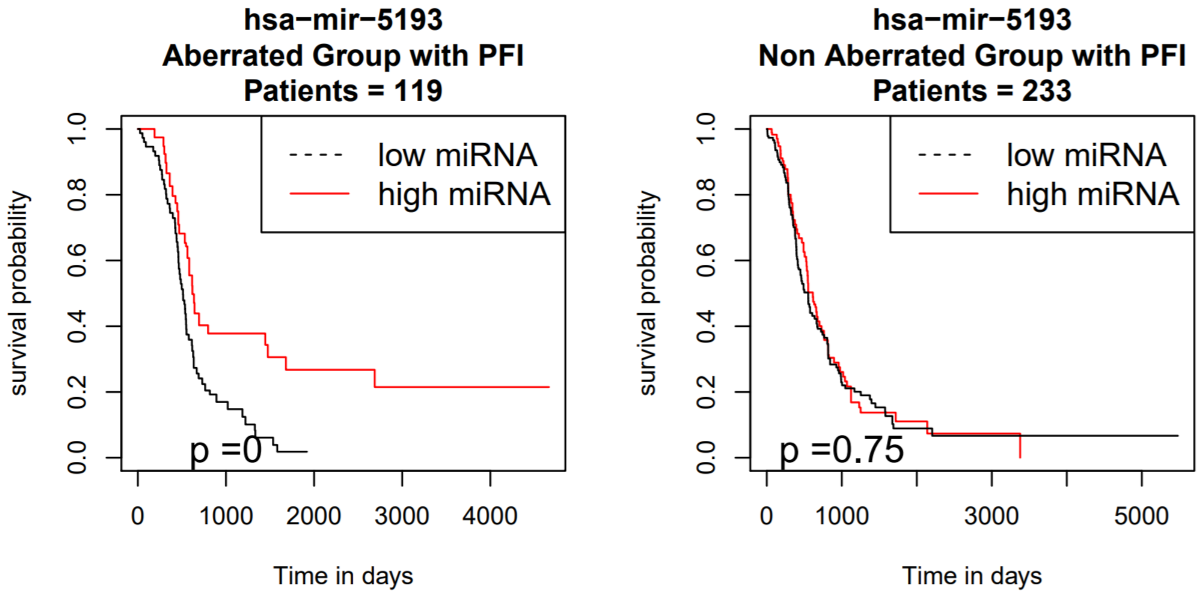

### Deletion Peak 16

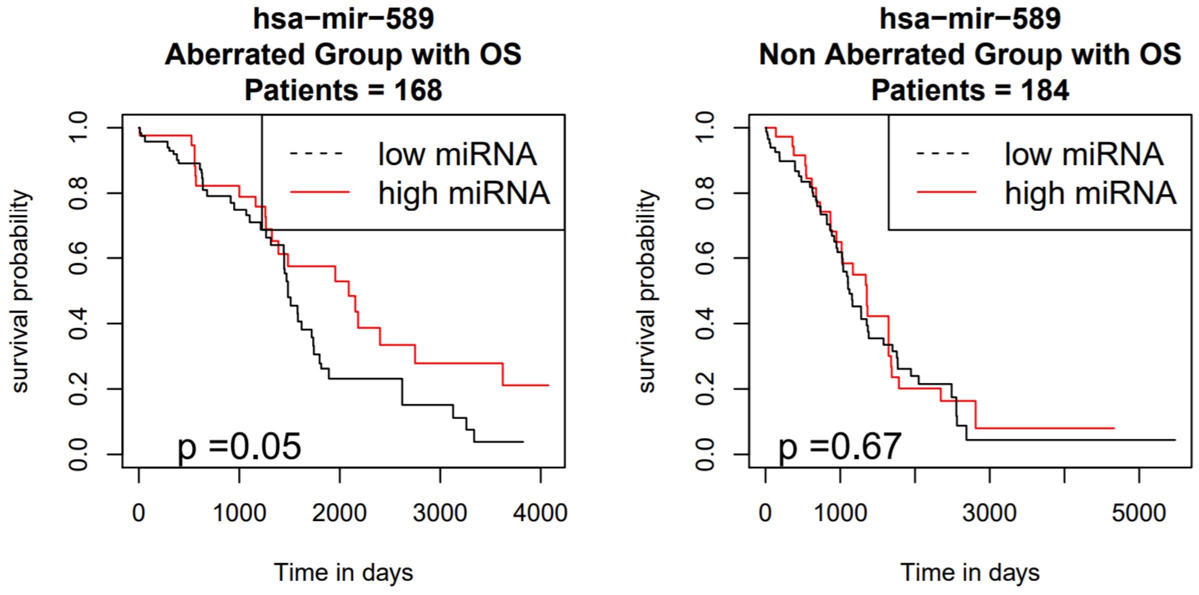


### Deletion Peak 16

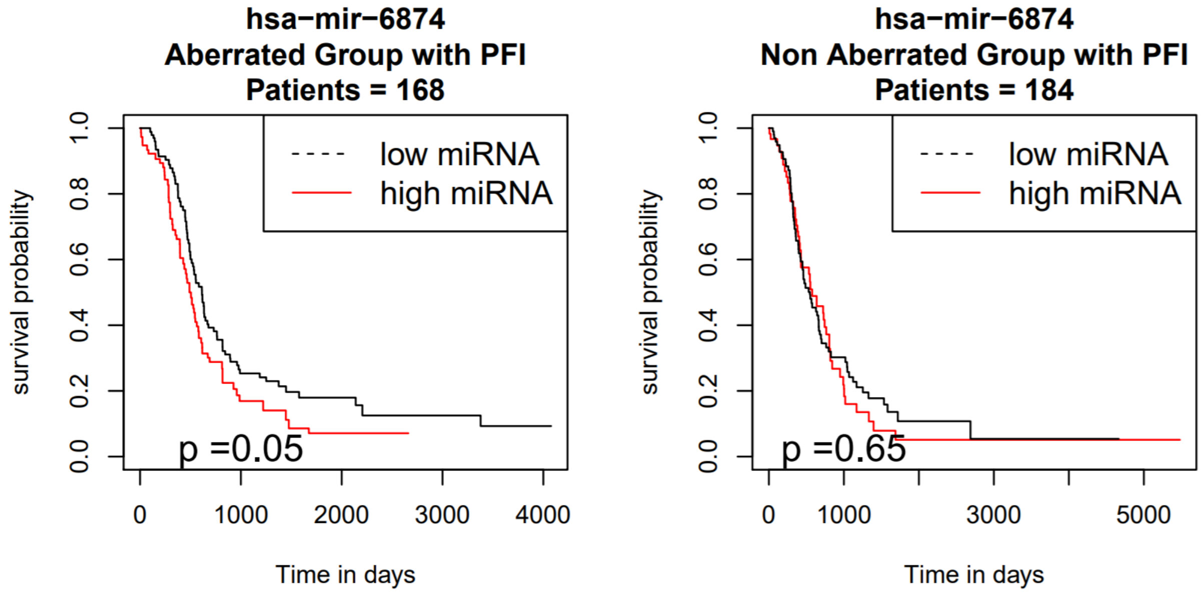

#### Deletion Peak 28

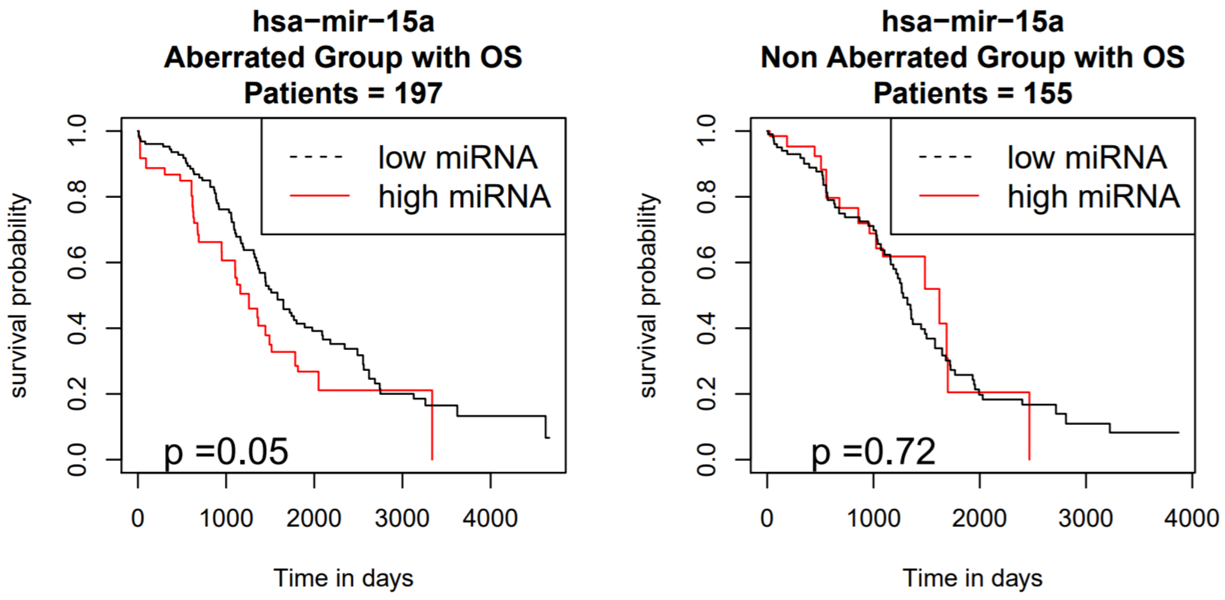


#### Deletion Peak 36

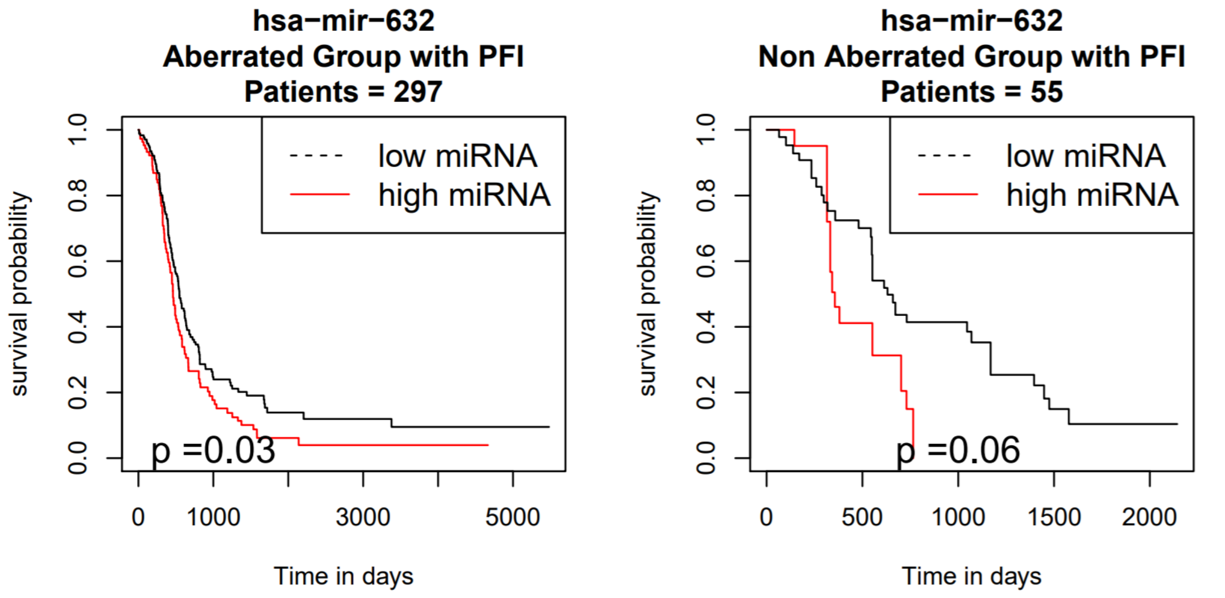

**Deletion Peak 43**

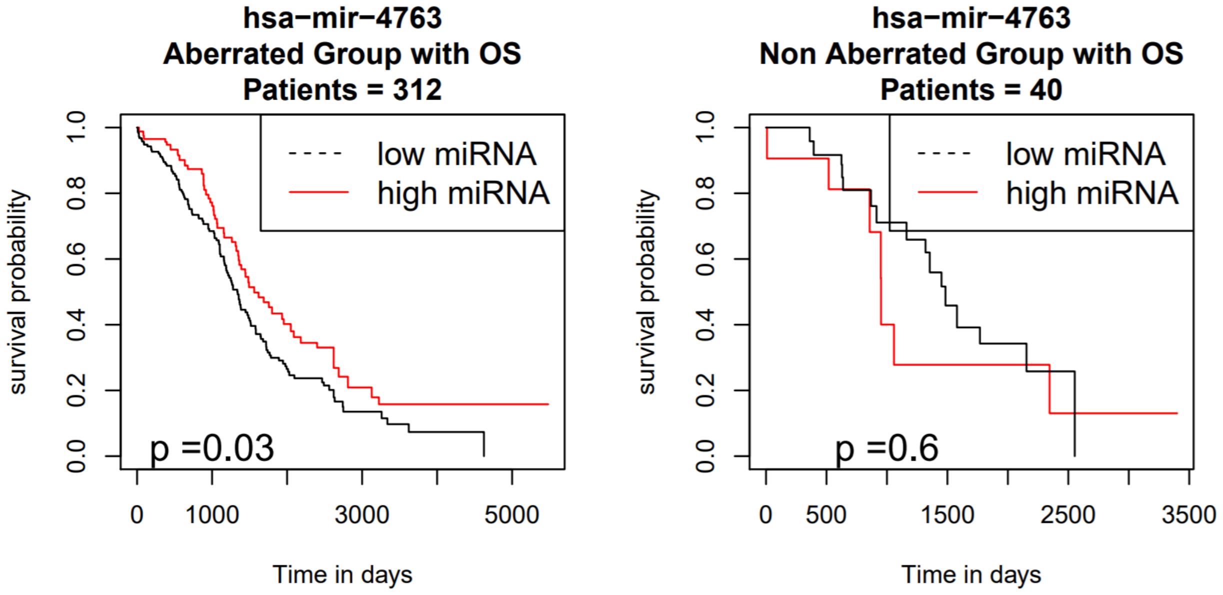
